# *In Vivo* Evolution of Monoclonal Antibody CR3022 to Achieve Cross-Neutralization of SARS-CoV-2 and Implications for Vaccine Strategies Against SARS-related Viruses

**DOI:** 10.1101/2025.08.05.666673

**Authors:** Yanbin Fu, Ziqi Feng, Steven A. Erickson, Peter J. Halfmann, Lei Li, Jordan C. Chervin, Chloe A. Troxell, Jiayi Sun, Atsuhiro Yasuhara, Siriruk Changrob, Min Huang, Nai-Ying Zheng, Meng Yuan, Yoshihiro Kawaoka, Ian A. Wilson, Patrick C. Wilson

## Abstract

The epitope that monoclonal CR3022 binds to represents a promising target for broad protection against a wide range of human and zoonotic coronaviruses. We developed a powerful model to evaluate antibody affinity maturation *in vivo* using immunoglobulin (Ig)-humanized mice that express the predicted germline heavy chain of antibody CR3022. SARS-CoV/SARS-CoV-2 sequential immunization led to the convergent evolution of the germline CR3022 through somatic hypermutation (SHM) that resembled the affinity-matured CR3022 from a human, but now also adapted to key variants and divergent sarbecoviruses. While simple prime-boost strategies drove CR3022-epitope targeting, an intensive vaccination protocol elicited dominant responses to other epitopes. X-ray crystal structures revealed that SARS-CoV-2-neutralizing CR3022-like antibodies exhibit enhanced affinity by increasing polar and electrostatic interactions. Overall, these findings show CR3022-like clones can be readily adapted through SHM to increase breadth and potency to sarbecoviruses by relatively minor shifts in affinity with appropriate vaccination strategies.

## Introduction

Betacoronaviruses include five viruses of great clinical importance to humans. Two of these, SARS-CoV and SARS-CoV-2 (the causative agent of COVID-19), are phylogenetically close and both are classified under the species SARS-related coronavirus ^1, 2, 3, 4^. Although the World Health Organization (WHO) declared an end to the global Public Health Emergency for COVID-19 in May 2023, the challenge posed by the virus remains significant, not only for immunocompromised populations but also for healthy individuals, as SARS-CoV-2 continues to evolve into increasingly infectious variants with higher immune escape potential ^5, 6, 7, 8, 9^. Furthermore, many zoonotic coronavirus strains found in animals pose a threat of future zoonosis events with pandemic potential ^10^. Broadly neutralizing antibodies (bnAbs) that target conserved sites within the RBD of the spike protein on the surface of SARS-CoV-2 virion offer potential for maintaining a broad spectrum of neutralizing activity against constantly evolving variants of concern (VOCs) and other SARS-related viruses within the sarbecovirus subgenus ^11, 12, 13, 14, 15, 16, 17, 18^. Among the conserved epitopes across sarbecoviruses, the site targeted by the monoclonal antibody (mAb) CR3022 was one of the earliest identified as a candidate for therapeutic intervention during the onset of the COVID-19 pandemic ^19, 20^. Originally isolated from a convalescent patient previously infected with SARS-CoV ^21^, CR3022 is cross-reactive to the RBDs of both SARS-CoV and SARS-CoV-2. However, it only demonstrated neutralizing potency against SARS-CoV, as its binding affinity to SARS-CoV-2 RBD was reduced by approximately 100-fold, although one study reported a measurable neutralizing activity for CR3022 against SARS-CoV-2 ^20, 22, 23, 24, 25, 26^. Previous studies have sought to enhance its binding affinity for SARS-CoV-2 RBD, either through *in vitro* site-directed mutagenesis or computational approaches ^20, 27, 28, 29, 30, 31, 32^. Unlike most RBD-targeting neutralizing antibodies, whose neutralization mechanism involves binding to the receptor-binding site (RBS) and direct competition with the cell receptor angiotensin-converting enzyme 2 (ACE2), CR3022 binds to a cryptic epitope outside the ACE2 binding site ^20, 33^. Structural studies have shown that CR3022 binding can induce conformational changes in the S1 domain of the spike protein, leading to dissociation of the trimeric spike ^24, 25^. Although CR3022 exhibits limited *in vitro* neutralizing activity against SARS-CoV-2, it has been shown to elicit diverse Fc effector-mediated immune responses, such as antibody-dependent cellular phagocytosis (ADCP) and antibody-dependent complement deposition (ADCD), even in the presence of ACE2 ^34^.

Here, we explored the potential of CR3022 to undergo affinity maturation *in vivo* using an Ig-humanized mouse model (glCR3022H) that expressed the predicted germline heavy chain of CR3022. To achieve a human-like, physiologically relevant CR3022-class B cell precursor frequency, we performed adoptive transfer of glCR3022H B cells into congenic recipients. We then assessed their ability to respond *in vivo* to priming immunization with soluble spike protein derived from either SARS-CoV or SARS-CoV-2. We demonstrated that germline CR3022 (glCR3022) binds to SARS-CoV but not to SARS-CoV-2, unless it undergoes affinity maturation toward the latter. Thus, whereas low-affinity SARS-CoV-2 spike immunogens failed to activate and recruit CR3022 precursors into germinal center (GC) reactions, a SARS-CoV/SARS-CoV-2 prime-boost sequential immunization regimen drove a cross-reacting response. This strategy allowed us to determine the affinity limits exerted by *bona fide* SHM targeting on the CR3022-like antibodies to SARS-CoV-2 variants. We identified novel, affinity-enhanced CR3022 variant clonotypes with on-track mutations resulting in new electrostatic and polar interactions that could adapt CR3022 antibody precursors to SARS-CoV-2 and variants, and to other sarbecovirus strains. Thus, this mouse model provides an excellent system for studying the biology of GC reactions and affinity maturation to an epitope on a complex and relevant protein antigen, with predicted IgV gene mutational outcomes converging between the mice and the original human CR30222 antibody. We also noted that more substantial immunization regimens, achieved by escalating dosage immunization ^35^, drove an almost complete replacement of the predominant CR3022-like B cells with endogenous B cells targeting other epitopes. This observation demonstrates that more robust immune stimulation protocols is not always beneficial, highlighting the complexity of targeting specific epitopes in the context of mechanisms such as serum-driven epitope masking or immunosubdominance. Our findings show that the CR3022 epitope is a readily targetable site of vulnerability *in vivo*, inducing cross-neutralizing antibody responses against various SARS-related viruses.

## Results

### Generation of a humanized Ig mouse model expressing predicted germline heavy chain of monoclonal antibody CR3022

To investigate the potential of mAb CR3022 for adaptation to acquire potent neutralizing activity against SARS-CoV-2 and related VOCs, we generated a transgenic glCR3022H mouse model. B cells from this model carry a rearranged variable region encoding the predicted germline heavy chain of mAb CR3022, comprising the human *IGHV5-51*\*03*, IGHD3-10\**01, and *IGHJ6*\*02 gene segments. The rearranged VDJ segments, fused with a murine *Ighv1-18* promoter, was integrated into the *Igh* gene locus via a CRISPR-Cas9-based system ^36^, and is therefore susceptible to natural SHM or class switch recombination (Fig. 1A). The CRISPR-Cas9 gene editing resulted in the deletion of varying numbers of nucleotides at the 3’ terminus of framework region 4 in all founders of the transgenic line for unknown reasons. Only one of these founders had an in-frame antibody gene but with a three-nucleotide deletion, resulting in the loss of the C-terminal serine residue. This serine deletion did not affect the binding, specificity, or expression of the glCR3022 antibody. Furthermore, the glCR3022H insertion functioned normally, and the mice readily expressed affinity-matured and class-switched CR3022-like heavy chains. The *Igl* gene locus was left intact to preserve combinatorial diversity. Genotyping confirmed that the native murine *Ighj* gene segments were deleted due to the integration of the targeting construct, thereby preventing most endogenous Ig rearrangements on the targeted allele (Supplementary Fig.1A, Supplementary Table 1). Unexpectedly, a natural cryptic recombination signal sequence (cRSS) was embedded within the CDRH3 region of the glCR3022H gene, as identified by an online prediction tool ^37^ (Supplementary Fig.1B). This cRSS caused the infrequent occurrence of hybrid V regions where the knock-in J gene segments rearrange with murine endogenous *Ighv* genes, resulting in distinct CDRH3 peptides in approximately 6% of the B cells (Data not shown). To determine the frequency of B cells expressing the glCR3022H knock-in sequence, we used flow cytometry to isolate single B220^+^/IgM^+^ splenic B cells from naïve glCR3022H mice and Sanger-sequenced the *Igv* genes after RT-PCR. The results showed that 90.6% of naïve B cells in heterozygous mice expressed the knock-in glCR3022H transgene (Fig. 1B). A comparison of the murine naïve B cell light chain repertoires between glCR3022H and wild-type (WT) mice revealed that a variety of light chain genes (mainly kappa) were expressed in the knock-in mice at frequencies similar to those in WT mice ^38^, suggesting that the human glCR3022H variable region could pair with various murine IgL rearrangements to form hybrid human-mouse BCRs (Fig. 1B and C) with a predominantly 9-residue CDRL3 (Fig. 1D).

**Figure 1.**
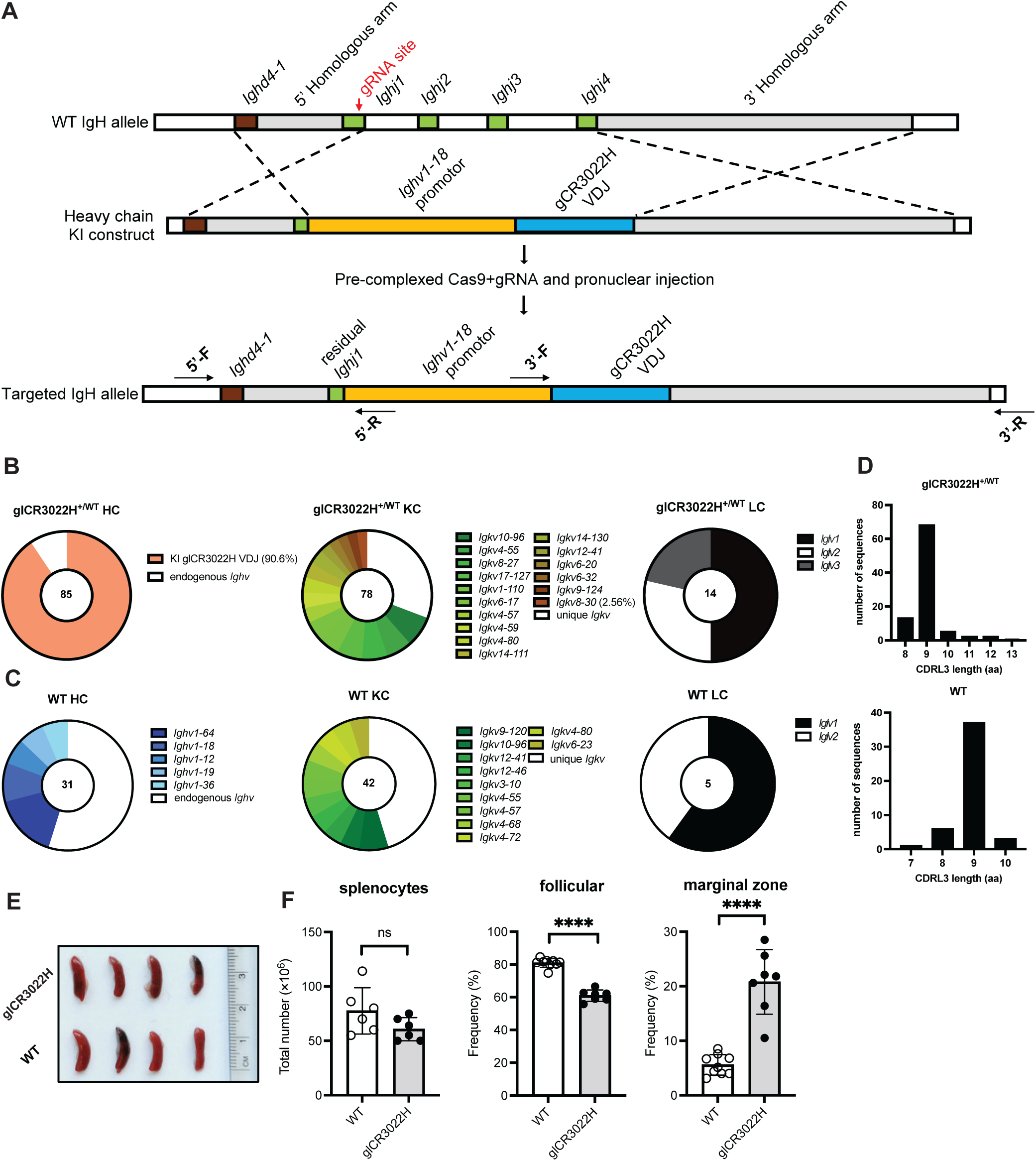
Generation and characterization of glCR3022H knock-in mouse. **(A)** Schematic figure of one-site cutting strategy with CRISPR-Cas9 method used for generating the glCR3022H mouse model. The homology regions are highlighted in light gray and indicated by dashed lines; The 5’ and 3’ primers used for genotyping knock-in founders and the descendants are shown with black arrows in the lower panel. **(B-C)** Pie charts show the V gene usage for heavy and light chain sequences amplified from single IgM^+^ splenic B cells sorted from unimmunized glCR3022H and control B6 mice. The number located in the center of the pie charts indicates the total number of sequences amplified. Any V genes found to be used by more than one sequence were shown with colored slices, and its size is proportional to the number of sequences using the V gene. The white slice indicates the proportion of unique V genes used. **(D)** CDRL3 lengths (aa) from light chains analyzed in (B) and (C). **(E)** Spleen size comparison between glCR3022H and B6 WT mice under unimmunized condition. **(F)** Absolute number of splenocytes and frequencies of follicular B cell and marginal zone B cell populations in the spleens of glCR3022H and control B6 mice are shown. The representative flow plots showing gating strategy for above B cell populations are depicted in supplementary Figure 1. \**p*<0.05, \*\**p*<0.01, \*\*\**p*<0.001, \*\*\*\**p*<0.0001(two-tailed Student’s *t* test). Each dot represents one mouse. Error bars indicate mean ± SD from mice in each group.

Spleen size and the absolute number of splenocytes in glCR3022H mice were comparable to those in WT C57BL/6 mice (Fig. 1E and F). However, as commonly observed in BCR knock-in models, there was a shift in the frequency of marginal zone (MZ) B cells and follicular (FO) B cells ^39^. Specifically, glCR3022H mice exhibited a significantly higher frequency of MZ B cells and a lower frequency of FO B cells in the spleen, as determined by flow cytometry (Fig. 1F and Supplementary Fig.1C). We next assessed early B cell development in the bone marrow, using the nomenclature by *Hardy et al*. ^40, 41^, and examined B cell compartments in the peritoneal cavity (Supplementary Fig.1D-I). Consistent with the expected course of Ig heavy-chain gene rearrangement during pro-B cell development, we observed a reduced frequency of early pro-B cells in the glCR3022H mice (Supplementary Fig.1F). This finding suggests that pro-B cells in these mice were quickly progressing to later developmental stages rather than undergoing *de novo* gene rearrangement. Additionally, a retention of cells was observed at the pre-B cell stage (Supplementary Fig.1G). Peritoneal B cells displayed near-normal frequencies of B-2 and B-1 subpopulations (Supplementary Fig.1H-I).

Collectively, we established a humanized BCR heavy-chain knock-in mouse model (glCR3022H) that expresses the predicted germline heavy chain of mAb CR3022 in a significant proportion of peripheral B cells exhibiting the knock-in BCR.

### Hybrid BCR expression on the B cell surface and antibody response elicited by immunization with SARS-CoV spike protein in glCR3022H mice

To determine whether glCR3022H B cells express functional BCRs composed of glCR3022H heavy chains paired with mouse-native light chains on the cell surface, we prepared tetrameric RBD probes from SARS-CoV and SARS-CoV-2 to assess the binding specificity of splenic B cells from naïve glCR3022H heterozygous mice. Flow cytometry revealed that approximately 1.73% of splenic B220^+^ B cells from naïve glCR3022H mice bound to SARS-CoV RBD, while only 0.002% bound to SARS-CoV-2 RBD (Fig. 2A and 2B). In contrast, naïve WT C57BL/6 mice showed much lower binding frequencies to SARS-CoV (0.237%) and slightly higher to SARS-CoV-2 (0.137%), indicating that the glCR3022HC transgene enhanced the binding of naïve B cells to SARS-CoV RBD, but not to SARS-CoV-2 RBD (Fig. 2A and 2B). Among SARS-CoV RBD-binding B cells, approximately 83% were epitope-specific, as confirmed by competitive blocking with a CR3022 fragment antigen-binding (Fab) region for the epitope on the probe (Fig. 2B).

**Figure 2.**
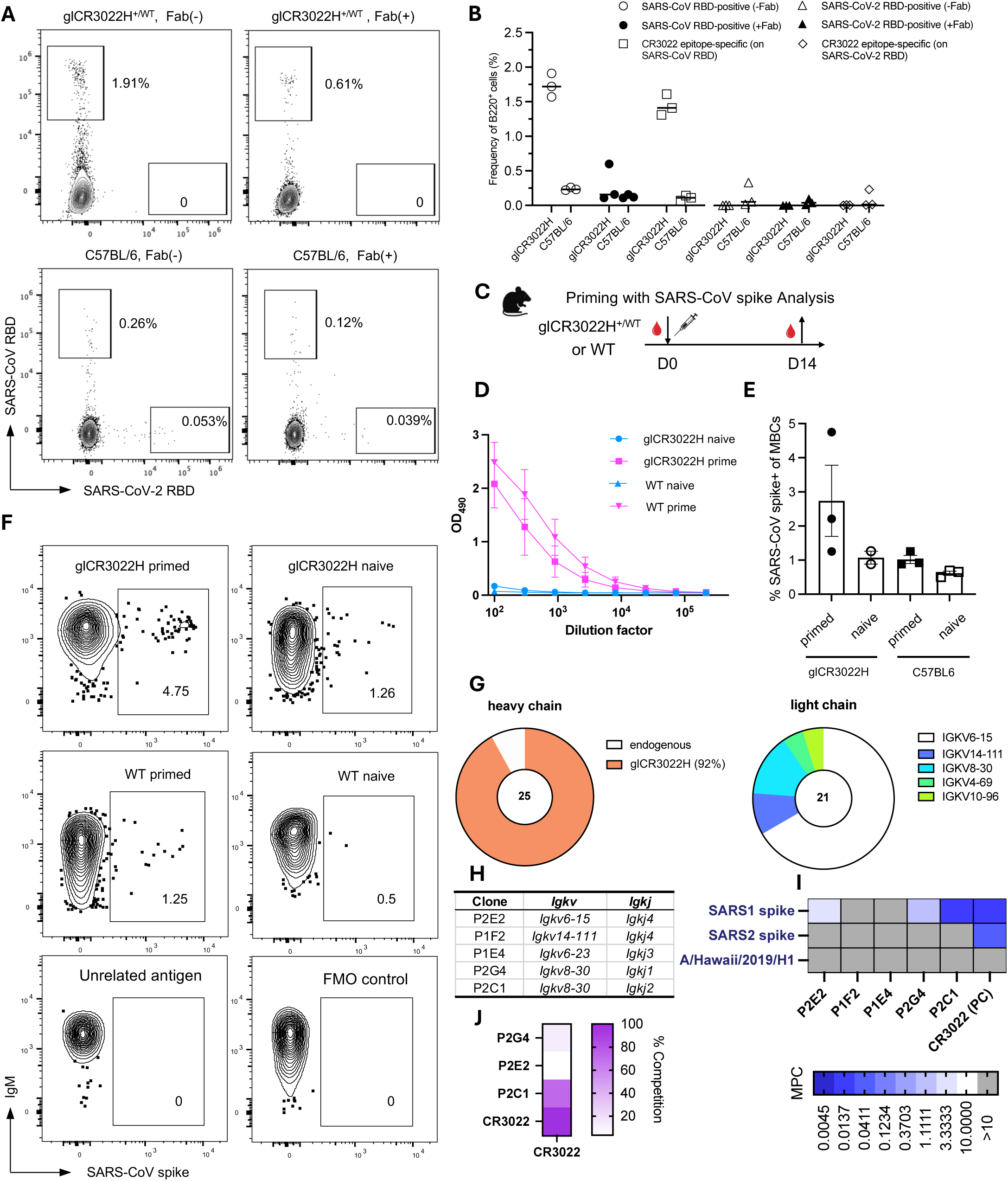
Hybrid BCR expression on B cell surface and antibody response elicited by immunization with SARS-CoV spike protein in gCR3022H mice. **(A)** Representative flow cytometry plots of CR3022’s epitope-specific B cells in naïve glCR3022H or control B6 mice. RBD from either SARS-CoV or SARS-CoV-2 were assembled as tetrameric probe with fluorescent streptavidin. Fab(+) group indicated that the probes were blocked in advance using an CR3022 Fab. B cells were gated on lymphocytes/singlets/viable/dump^-^/B220^+^ cells. **(B)** The proportion of CR3022’s epitope-specific B cells in naïve glCR3022H and B6 mice as shown in (A), n = 3. **(C)** Schematic of priming regimen with SARS-CoV spike protein. **(D)** Serum ELISA quantification of IgG titer against SARS-CoV spike for immunized or naive glCR3022H and B6 mice. Error bars are mean ± SD. **(E)** The quantification of proportion of immunogen-specific MBCs as shown in (F). Error bars are mean ± SEM. **(F)** Representative flow cytometry plots showing the frequency of immunogen-specific MBCs on day 14 post priming. MBC were gated on lymphocytes/singlets/viable/dump^-^/B220^+^/IgD^-^/CD38^+^/antigen^+^.**(G)** V genes usage of immunogen-specific MBCs single cell-sorted from (E). (H) *Igkv* and *Igkj* usage of clonally representative mAbs selected for verifying the binding specificities. **(I)** Heatmap of minimal positive concentration(MPC) of the mAbs tested for binding SARS-CoV or SARS-CoV-2 spike and an unrelated antigen, haemagglutinin (HA) from A/Hawaii/70/2019 (H1N1) strain. **(J)** Competition of SARS-CoV spike-binding mAbs with parental CR3022.

Following immunization with soluble SARS-CoV spike trimer formulated with the adjuvant poly(I:C) (Fig. 2C), anti-SARS-CoV spike IgG titers were detected in both knock-in and WT mice on day 14 post-immunization by ELISA (Fig. 2D). SARS-CoV spike-specific memory B cells (MBCs) were isolated from the spleens of glCR3022H mice at day 14 post-immunization by flow cytometry using a SARS-CoV spike probe (Fig. 2E and 2F) ^42^. Among all murine *Igkv* genes, *Igkv8-30*, with 84.1% nucleotide identity, is the most similar to human *IGKV4-1* that encodes the V region of the CR3022 light chain and is expressed at a frequency of 2.56% in B cells before immunization (Fig. 1B). BCR sequencing of single-cell sorted SARS-CoV spike-binding MBCs showed that 92% of the cells expressed the glCR3022H transgene, with a six-fold enrichment of *Igkv8-30* usage accounting for 14% of the Ig light chain gene repertoire (Fig. 2G). To validate the specificity of these MBC-derived BCRs, we cloned and expressed five recombinant monoclonal antibodies representing all the clonotypes from the sorted MBCs (Fig. 2H). As expected, only hybrid antibodies using mouse *Igkv8-30*, such as P2G4 and P2C1, were shown to bind the CR3022 epitope on SARS-CoV spike antigen, albeit with varying binding affinities. In contrast, antibodies expressed by glCR3022H with other murine *Igkv*, such as P2E2, P1F2, and P1E4, either bound to unknown epitopes (P2E2) or did not bind the spike antigen at all (Fig. 2I and 2J). For example, P2E2 used gCR3022 with *Igkv6-15*, and typified another clonal expansion that bound SARS-CoV spike, but generally with low affinity at a non-overlapping epitope that was non-neutralizing. Interestingly, antibodies P2C1 and P2G4 using *Igkv8-30*, which mainly differ in their usage of *Igkj* genes (Fig. 2H), exhibited a notable difference in binding affinity, with P2G4 showing a substantial reduced binding compared to P2C1 (Fig. 2I and Supplementary Fig.2A). Subsequently, 23 additional antibodies encoded by glCR3022H and *Igkv8-30* encoded light chains validated this observation on J chain usage (Supplementary Table 2). Moreover, the absence of CR3022 epitope binding in three pairings of glCR3022H with other light chains demonstrated that *Igkv8-30* was essential for binding to SARS-CoV spike in mice (Supplementary Table 2).

Next, we investigated the monovalent binding affinities of P2C1, reverted to its germline form, for SARS-CoV and SARS-CoV-2 RBDs, along with the germline and affinity-matured CR3022. We hypothesized that B cells bearing the germline-reverted P2C1 BCRs would resemble the naïve CR3022 precursors. Germline P2C1 Fab was expressed and measured for binding to SARS-CoV RBD, yielding an equilibrium dissociation constant (K_D_) of 558 nM that was 3.8-fold lower than that of glCR3022 Fab (Supplementary Fig.2B). In contrast, germline P2C1 Fab showed negligible binding to SARS-CoV-2 RBD, with binding below the limit of detection (Supplementary Fig.2B). Consistent with previous studies ^20, 23^, CR3022 Fab exhibited K_D_ of 7.3 nM for SARS-CoV RBD and 79.1 nM for SARS-CoV-2 RBD (Supplementary Fig.2B).

The loss of affinity by the original CR3022 heavy/light chain pairing without accumulated somatic mutations was predictive of a loss in neutralization capacity. Furthermore, the glCR3022H transgene paired with *Igkv8-30*-encoded light chains, which bind the CR3022 epitope, but with variant CDRL3 junctions, might compensate and allow neutralization without the need for somatic mutations. We therefore assessed the neutralizing activities of predicted germline-reverted P2C1 compared to the original CR3022 against SARS-CoV and SARS-CoV-2 pseudoviruses. Only affinity-matured CR3022 or transgene-encoded IgGs were able to neutralize the SARS-CoV pseudovirus (Supplementary Fig.2C), with comparable IC_50_ values, indicating a dependence on affinity maturation.

In summary, we demonstrated that the glCR3022H mouse can mount an immune response following SARS-CoV spike priming immunization, with B cells expressing knock-in heavy chains paired with murine *Igkv8-30* behaving similarly to CR3022. Germline-reverted CR3022-like precursors exhibit 290-fold reduced affinity for SARS-CoV RBD, and no binding to SARS-CoV-2 RBD, and do not neutralize SARS-CoV or SARS-CoV-2.

### CR3022-like B cells, present at physiological precursor frequencies, can be efficiently recruited into germinal centers following SARS-CoV but not SARS-CoV-2 spike priming

As CR3022 was initially isolated from a person infected with SARS-CoV and did not exhibit germline affinity to SARS-CoV-2 (Supplementary Fig.2B, C), we investigated whether priming with the SARS-CoV-2 spike could induce B cell responses. For this experiment, we used a congenic transfer model to assess the response with a normalized frequency of CR3022-like germline precursor cells. The precursor frequency of CR3022-like B cell clones in humans was predicted using publicly available BCR repertoire datasets generated by single-cell RNA sequencing (scRNA-seq), which contain paired heavy and light chain configurations ^43^. CR3022 was originally isolated as a single-chain variable antibody fragment (scFv) from independent libraries of *IGHV* and *IGLV* genes from a SARS-CoV convalescent patient ^26^. Therefore, the V_H_ and V_L_ of CR3022 may be derived from two unrelated antibodies. We were interested in identifying any naturally occurring clones similar to CR3022 in humans. Filtering antibody candidates using stringent public clone-defining criteria from the CoV-AbDab database ^44^, we did not identify any clones with identical *IGV* and *IGJ* gene usages as CR3022. However, several clones with similar V_H_ and V_L_ pairings were identified that had binding and neutralizing characteristics like CR3022. For example, one such clone, COVA3-10, which differs only in the *IGKJ* gene usage, bound both SARS-CoV and SARS-CoV-2 RBDs, albeit with lower affinity, and did not compete with ACE2 for RBD binding ^45, 46^. These features suggest that COVA3-10 belongs to the category of cross-reactive mAbs targeting the typical class 4 epitope on RBD of SARS-CoV-2, similar to CR3022 ^47, 48^. Precursors similar to CR3022, using *IGHV5-51*/*IGHJ6* heavy chains and *IGKV4-1*/*IGKJ2* light chains, occurred at a frequency of approximately 11 per 10^5^ naïve B cells across data from four human donors in the database (Supplementary Fig.2D). B cells using these VJ pairings and the *IGHD3-10* and *IGHD5-18* genes, encoding partial CDRH3 sequences of CR3022 and COVA3-10, respectively, were enriched in CR3022-like clones identified from the B cell compartments of two donors with a history of SARS-CoV-2 exposure (Supplementary Fig.2E). These findings suggest that CR3022-like precursors likely exist and can be activated upon natural SARS-CoV-2 infection. Based on these analyses, the frequency of transgenic CR3022-like precursor B cells paired with murine *Igkv8-30* was adjusted to approximate the physiological precursor frequency estimated for humans (Supplementary Fig.2D). For preliminary analysis, 10^6^ donor naïve B cells enriched from the glCR3022H transgenic mice that express the congenic marker CD45.2 were transferred into recipient B6 mice (CD45.1^+^), resulting in 79 CD45.2 donor B cells out of 122,311 B220^+^ B cells in the recipient spleen 3 to 4 days after transfer (Fig. 3A). With a goal reflective of the 1.5% of naïve human B cells that are CR3022-like precursors (Fig. 2B), and considering that 2.5% of transgenic glCR3022H B cells pair with the *Igkv8-30* light chain before immunization, we estimated that the transfer of 5 million total transgenic B cells in subsequent assays would achieve precursor frequencies within the physiological range of 1 to 5 CR3022-like cells per 10^5^ total B cells (Supplementary Fig.2D).

**Figure 3.**
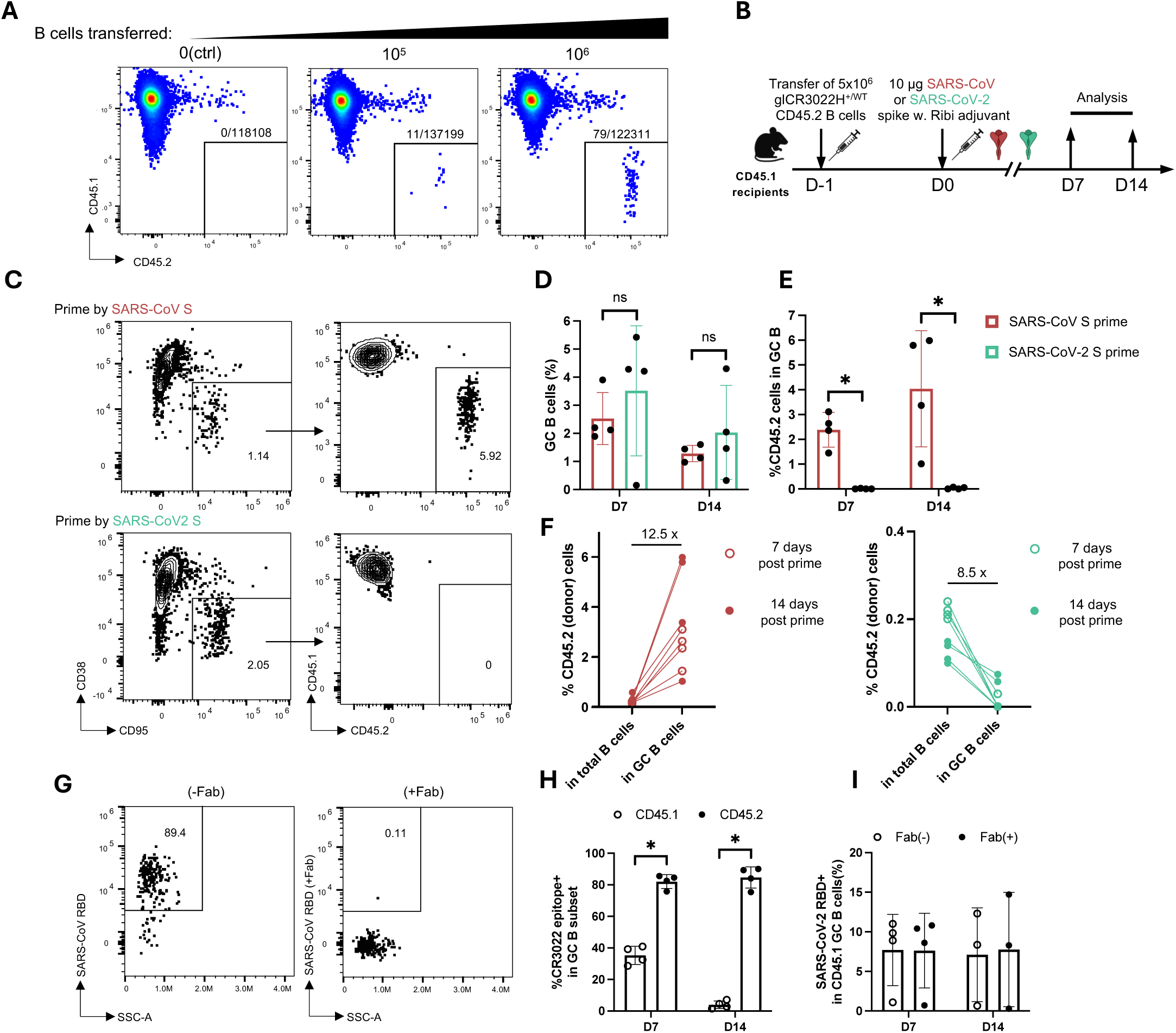
CR3022-like B cells at physiological precursor frequency can be efficiently recruited into germinal centers upon SARS-CoV spike prime. **(A)** Representative flow plots showing the proportion of CD45.2 donor cells detected in peripheral B cells of CD45.1 recipient mice was positively proportional to the quantities of CD45.2 donor cells transferred in; **(B)** Schematic of adoptive transfer conducted at CR3022-like precursor frequencies of ∼5 per 10^5^ B cells on one day prior to priming immunization with SARS-CoV or SARS-CoV-2 spike adjuvanted by Ribi. Samples were collected on day 7 and 14 post immunization for flow analysis; **(C)** Representative flow plots showing GC B cell proportion among B220^+^ B cells and frequencies of CD45.2 B cells recruited into GC compartment between two immunization groups on the day 14 post the primes; **(D-E)** Quantitative results for panel C (n=4 mice per group and per time point, mean±SD); **(F)** The comparison of the frequencies of CD45.2 B cells among total B cells between that in GC B cells subsets for two immunization groups on two time points; **(G)** Representative flow plots showing the percentage of SARS-CoV RBD probe^+^ B cells significantly decreased when a Fab-blocking probe was used for determining the epitope-specific GC B cells; **(H)** Quantitative results demonstrating that for SARS-CoV spike-primed group, the percentage of CR3022 epitope-reactive B cells in CD45.1 or CD45.2 GC B cell compartment on each of the time points; **(I)** Quantitative results showing that for SARS-CoV-2 spike-primed group, the percentages of SARS-CoV-2 RBD reactive-B cells in CD45.1 GC B cell compartment when two forms of probes were used. Error bars in all quantification plots indicate mean ± SD, \**p*<0.05 (two-tailed Student’s *t* test). Each dot represents one mouse.

After transfer, we analyzed the immune responses at days 7 and 14 post-priming with SARS-CoV or SARS-CoV-2 spike antigens formulated in Ribi oil-in-water emulsion adjuvant (Fig. 3B). At both time points, the GC responses were comparable between the two groups (Fig. 3C-D), though GC reactions waned over time. In SARS-CoV–primed recipient mice, there was a marked enrichment of CD45.2⁺ donor-derived cells within the GC response, exhibiting an average 12.5-fold increase in their proportion relative to the total B220⁺ B cell population (Fig. 3E-F). In contrast, in SARS-CoV-2-primed mice, almost all GC B cells were of recipient (CD45.1^+^) origin, indicating that CR3022-like precursors from the transgenic donor mice were unable to engage the SARS-CoV-2 antigen and enter the GC (Fig. 3C, E, F). The frequency of donor cells in the GC compartment of SARS-CoV-2-primed mice was even lower than their frequency in the total B cell population (Fig. 3F, right panel). This result is consistent with our expectation, as the predicted germline CR3022-like Fab exhibits negligible affinity for the SARS-CoV-2 RBD but retains substantially higher affinity (∼550 nM) for the SARS-CoV RBD. Using tetrameric antigen probes conjugated with fluorescently labeled streptavidin, we found that 80-90% of CD45.2 GC B cells in SARS-CoV-primed mice were CR3022 epitope-specific, as confirmed by a Fab-blocking probe encoded by CR3022 (Fig. 3G-H), which represents a 53-fold increase in the proportion of CR3022-epitope-specific B cells (80% vs. 1.5%). However, no SARS-CoV-2 RBD-reactive cells were detected at this point (data not shown). Notably, at day 7 post-immunization, the binding of 30% – 40 % of non-transgenic (CD45.1^+^) GC B cells could be competitively inhibited by the CR3022-Fab, suggesting that B cells binding epitopes proximal to the CR3022 epitope were readily induced during the early stages. However, by day 14, the proportion of endogenous GC B cells binding epitopes overlapping with CR3022 decreased to approximately 4%, implying that other GC clones binding more dominant epitopes rapidly outcompeted these cells as the GC reaction progressed. In SARS-CoV-2-primed mice, the proportion of SARS-CoV-2 RBD^+^ cells remained similar regardless of whether an Fab-free or Fab-blocking probe was used (Fig. 3I). This suggests that the B6 mice lack B cells binding the CR3022 epitope on SARS-CoV-2 RBD, which is likely due to the presence of four residue differences between SARS-CoV and SARS-CoV-2 at this epitope ^23^.

In summary, our data here demonstrate that, when starting with a physiological precursor frequency, CR3022-like precursors from glCR3022H mice can be efficiently activated by SARS-CoV spike priming. These data also suggest that targeting of the conserved CR3022 epitope on SARS-CoV-2 requires priming first with SARS-CoV. Furthermore, it appears that endogenous GC B cells reacting to epitopes proximal to the CR3022 epitope were readily induced in GCs early on but rapidly declined from 30% to 4% between days 7 and 14. This observation provides some insight into potential mechanisms of immunodominance, suggesting that, even if readily targeted by the base repertoire, epitopes more easily adapted to become dominant in GCs over time.

### Single-cell RNA-seq revealed that different immunization regimens directed distinct immune responses

It remains unclear how CR3022-like antibodies can be adapted for neutralizing SARS-CoV-2 and its variants *in vivo*. To explore how SARS-CoV-2 spike-adapted CR3022 variants might evolve potential broad neutralizing capabilities, we used and compared three sequential immunization strategies in glCR3022H transgenic mice (Fig. 4A). All three strategies were primed with the SARS-CoV spike and then boosted with SARS-CoV-2 spike or spikes of relevant variants, each formulated in Ribi adjuvant, and for at least two doses. In Cohort 1, mice were sub-grouped into two groups: one was boosted with ancestral (Wuhan strain) SARS-CoV-2 spike (n=12), and the other were boosted with omicron spikes (XBB.1, BQ.1, or XBB.1.5) (n=12). Mice in cohort 2 (n=6) received an additional boost dose to isolate plasmablasts (PBs) that are thought to arise from reactivated MBCs that peak about a week post-vaccination ^49^. Cohort 3 (n=8) was subjected to an escalating dose regimen, as previous studies have shown that such protocols can enhance humoral responses to subdominant epitopes, particularly in models of HIV antigen immunization ^35, 50^. At the end of each immunization course, we harvested spleens and mesenteric lymph nodes (mLNs) from all cohorts, isolating and immunomagnetically enriching pan-B cells (Supplementary Fig.3A). Antigen-specific B cells were baited using fluorescently labeled streptavidin conjugated to biotinylated SARS-CoV-2 RBD and oligonucleotide tags (Supplementary Fig.3A-B). For PB isolation, a bulk-sorting strategy was employed due to the low BCR density on the surfaces of antibody-secreting cells (ASCs) that precluded detection with fluorescent probes (Supplementary Fig.3C) ^51, 52^. We then performed scRNA-seq using the 10x-Genomics platform to obtain transcriptomic and paired mouse BCR V(D)J repertoire data from 38 mice across all immunization groups.

**Figure 4.**
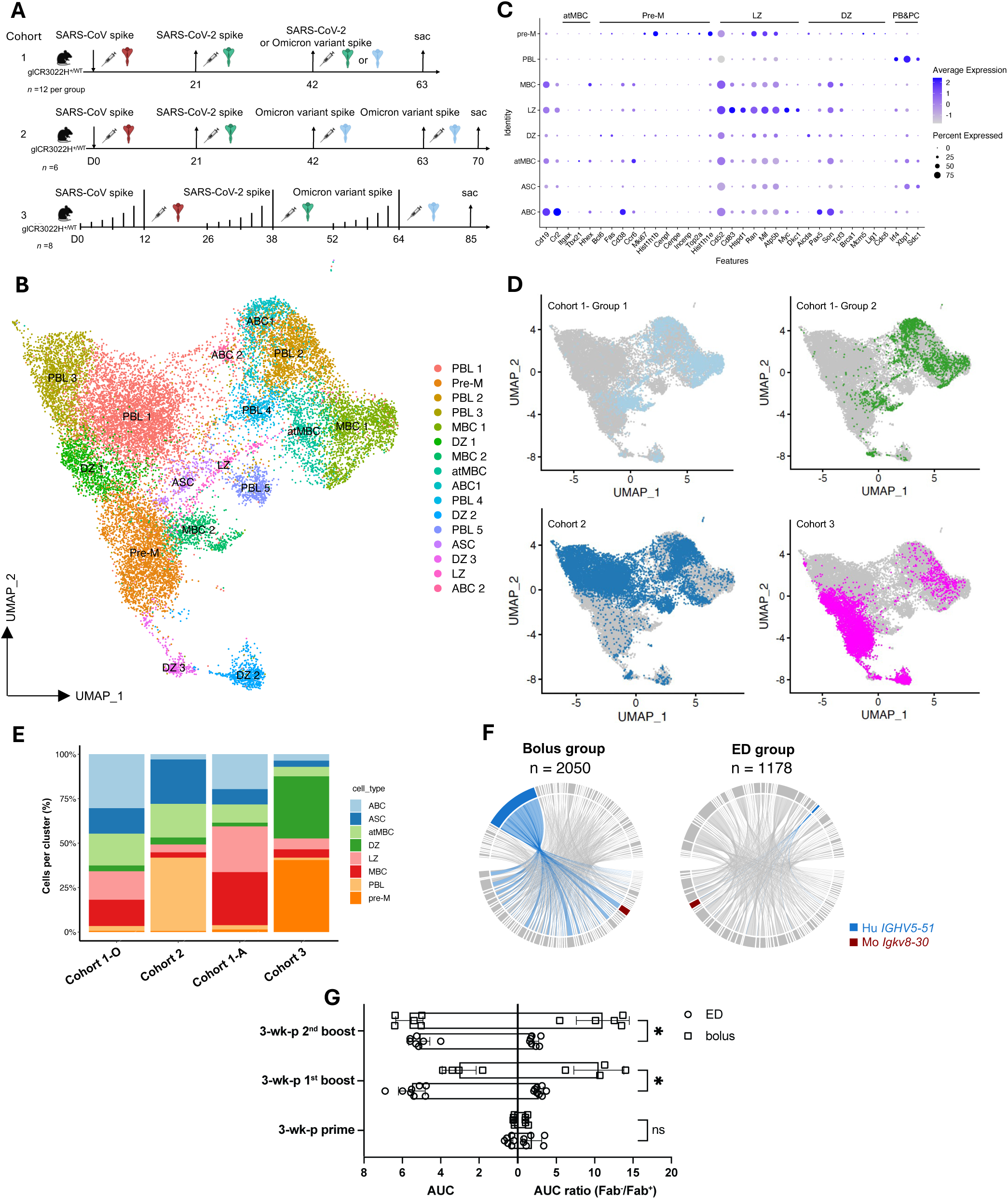
Single-cell RNA-seq revealed different immunization regimens directed distinct immune response. **(A)** Schematic of three immunization regimens tested on glCR3022H mice; **(B)** Uniform Manifold Approximation and Projection (UMAP) of antigen-specific glCR3022H B cells and bulk-sorted plasmablasts pooled from all mice. Each cell is represented by a dot and colored by the cluster it is belonging to; **(C)** Dot plot of gene expression for representative gene markers used for annotating the identities of each B cell subset; **(D)** UMAP showing the distribution of B cells from each of the immunization groups. Group1-A and group 1-O in the upper panels indicate the cells from subgroup of group1 where the WT (Wuhan) strain of SARS-CoV-2 spike was used in the last boost and where Omicron spike was used, respectively; **(E)** Bar graph breaking down the B cells compositions of each cohort into discrete cell clusters; **(F)** Comparison of percent of CR3022-like clones between bolus and escalating dosing groups, pairing of heavy and light chains were shown by circos plot; **(G)** Mouse anti-SARS-CoV-2 RBD titer (AUC) and AUC ratio (ratio of AUC in the absence of Fab to AUC in the presence of Fab) is shown for cohort 2 (bolus dosage) and cohort 3 (escalating dosage). Error bars in all quantification plots indicate mean ± SD, \**p*<0.05 (Mann-Whitney U tests). Each dot represents one mouse.

From the transcriptomes of 20,171 sequenced cells, which included both antigen-specific B cells and plasmablasts, we identified 16 distinct transcriptional clusters (Fig. 4B). Gene expression analysis and Gene Set Enrichment Analysis (GSEA) revealed six major B cell subsets: plasmablast-like (PBL), activated B cells (ABC), MBC, atypical memory B cells (atMBC), ASCs, and GC B cells. GC B cells were further subdivided into three sub-clusters: dark zone (DZ) B cells, light zone (LZ) B cells, and pre-memory (Pre-M) cells (Fig. 4B-C). PBL and ASCs were identified by expression of high levels of *Irf4*, *Xbp1*, and *Sdc1* (CD138); while DZ cells are marked by *Aicda*, *Pax5*, and *Tcf3* ^53^; and *Cd83*, *Cd52*, and *Mif* distinguishable LZ cells ^54^. Pre-M cells, as a transitional population, were characterized by high expression of *Hist1h1b*, *Hist1h1e*, and *Mki67*^55^. Typically, more than one cluster can be identified that expresses genes associated with a particular B cell subset, we therefore assigned a stochastic number behind the clusters to distinguish sub-clusters (e.g., PBL1, PBL2, etc.). As expected, antigen-specific cells from all three immunization groups were predominantly composed of ABCs, MBCs, atMBCs, GC B cells, and PBLs (Fig. 4D, upper panels). Consistent with expectations, immunization Cohort 2, which included a bulk-sort of PBs, had a much higher proportion of cells clustered as PBLs and ASCs in the UMAP analysis (Fig. 4D-E). Escalating dosage immunization (Cohort 3) had a strong enrichment for GC B cells, with approximately 80% of cells exhibiting features of GC B cells (Fig. 4E). These results suggest that the escalating dosing regimen effectively promoted GC B cell differentiation and expansion, consistent with findings from HIV immunization studies in rodents and nonhuman primates ^35, 50^. Furthermore, different immunization regimens can yield distinct profiles of B cell subsets and also highlight the heterogeneity of B cells within the SARS-CoV-2 RBD^+^ population.

To assess the dynamics of CR3022-like B cell induction, we analyzed the BCR repertoire of 3,228 sorted B cells. Notably, the proportion of B cells with the knock-in heavy chain paired with murine *Igkv8-30* dropped significantly across all immunization groups. We hypothesized that this reduction occurred because the CR3022 epitope, although conserved, is a relatively inaccessible and cryptic antigenic site on spike RBD ^20^, likely outcompeted by dominant immune responses against more immunodominant and exposed epitopes (e.g., the RBS on RBD) ^20^. This drop in CR3022-like B cell induction was particularly evident in the escalating dose group (Fig. 4F). Consistent with the BCR repertoire analysis, serum titers from bolus dosing groups 1 and 2 showed a stronger response to the CR3022 epitope on the SARS-CoV-2 RBD (Fig. 4G). In contrast, serum from the escalating dose group had a broader response to multiple RBD epitopes (Fig. 4G). This finding aligns with prior studies suggesting that slow-delivery immunization protocols tend to favor the induction of more diverse immune responses by modulating the immunodominance of non-neutralizing epitopes ^50^. However, unexpectedly, the escalating dosing strategy did not effectively improve the immune response to the less immunodominant CR3022 epitope in this model. These observations further support the notion that immunosubdominance is driven by a more ready adaptation to competing epitopes in GC responses.

### CR3022-like antibodies underwent convergent evolution during affinity maturation and were easily adapted for neutralizing SARS-CoV-2 and related VOCs

To thoroughly evaluate the CR3022-like antibody variants adapted in sequentially immunized transgenic mice, we expressed 23 mAbs representing B cell clones encoded by glCR3022H and *Igkv8-30*, originating from various B cell subsets (Supplementary Fig.4 A-B, Supplementary Table 2). These clones exhibited a broad range of hypermutation levels (Supplementary Fig.5A-C). To evaluate adaptation to SARS-CoV-2 and cross-reactivity with variants and other sarbecoviruses, we tested the binding of the 23 mAbs to a comprehensive panel of 11 different RBD and spike antigens (Fig. 5A-B). Notably, 87% (20 out of 23) of the antibody candidates displayed high binding activity to SARS-CoV-2 and VOCs, regardless of whether the omicron spike antigens were used as immunogens for the final boost. In contrast, the parental CR3022 antibody, while cross-reactive between SARS-CoV and SARS-CoV-2, showed substantially reduced binding to Omicron VOC antigens. A large portion of the adapted CR3022 variants retained medium to high reactivity toward spike proteins from more phylogenetically distant zoonotic strains BM48-31 (35%) and HKU3-8 (78%). These findings highlight the conserved nature of the CR3022 epitope across sarbecoviruses.

**Figure 5.**
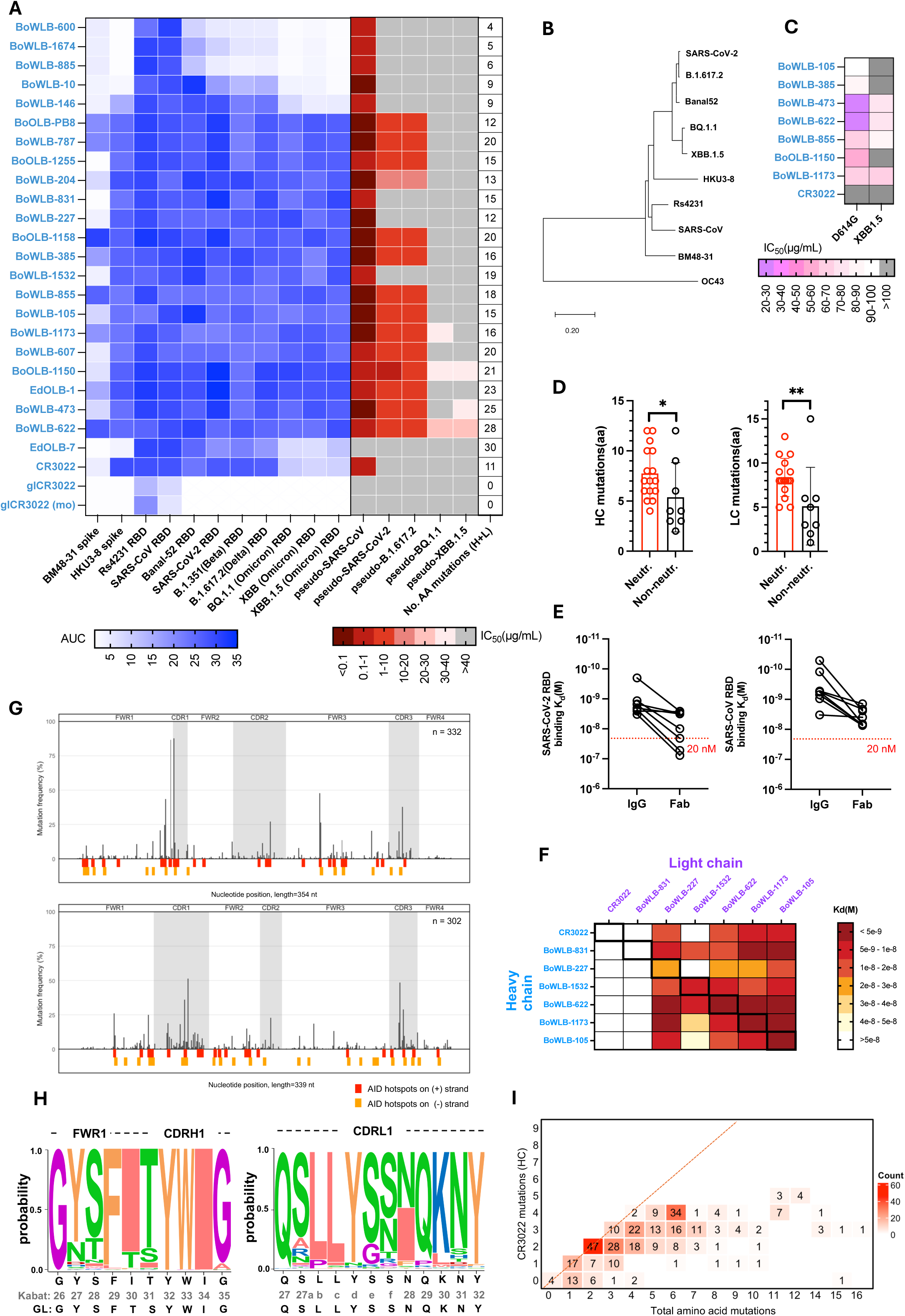
CR3022-like antibodies underwent strong selective pressure during the process of affinity maturation and were readily to be adapted to neutralize SARS-CoV-2 and related VOCs under proper immunization protocol. **(A)** Heatmap of binding spectrum toward spike or RBD antigens of different Sarbecoviruses, neutralization potency and number of total amino-acid mutations of expressed 23 mAbs. IDs for mAbs begin with two letters denoting the immunization strategy (Ed for escalating dosage, Bo for bolus), followed by three letters indicating the viral strain used as the final boosting immunogen (e.g., Wuhan, Omicron); **(B)** Phylogenetic tree of Sarbecoviruses based on the RBD sequences, the OC43 was used as an outgroup; **(C)** Heatmap of neutralization IC_50_ against SARS-CoV-2 D614G and XBB.1.5 variant; **(D)** Comparison of total amino-acid mutation numbers between neutralizing and non-neutralizing CR3022-like mAbs; **(E)** Change of binding affinity constants when CR3022-like mAbs were expressed as Fab or bivalent IgG; **(F)** Shuffled heavy (rows) and light chains (columns) from CR3022-like antibodies were tested for binding affinity to SARS-CoV-2 RBD; **(G)** Manhattan plots showing the mutation frequency in the HC (top panel) or LC (bottom panel) nucleotide sequences of CR3022-like antibodies pooled from all groups of immunizations; **(H)** Weblogo plot showing the residue composition on each specific position for CR3022-like HC and LC sequences; **(I)** CR3022 mutations detected in the knockin *IGHV5-51* genes. The y-axis indicates the number of CR3022 mutations observed in a given CR3022-like variant adapted in transgenic mice. The number within each square on the plot represents the number of antibody HC sequences that contained those specific numbers of CR3022-type mutations. The x-axis indicates the number of mutations that a set of CR3022-like HC sequences accumulated.

Consistent with prior reports^20, 22, 23, 56, 57^, the parental CR3022 antibody was ineffective at neutralizing SARS-CoV-2 and VOCs. However, approximately 60.9% (14/23) of the adapted CR3022 variants exhibited neutralizing activity against both SARS-CoV-2 (Wuhan) and B.1.617.2 (Delta) variants, while 13.0% (3/23) were able to neutralize the XBB.1.5 variant (Fig. 5A). Relative to the predicted binding affinity, a notable discrepancy was observed in neutralization potency for the Omicron variant pseudovirus assays. To verify Omicron neutralization, we tested seven antibodies with high binding affinity to Omicron variants, detectable pseudoviral neutralization, and full competitive inhibition of binding by CR3022 mAb using authentic live Omicron XBB.1.5 viruses (Fig. 5C). All seven antibodies neutralized the SARS-CoV-2 D614G variant at 20-100 μg/mL, and four cross-neutralized the Omicron XBB.1.5 variant virus with generally reduced IC_50_, despite no prior exposure to this variant. It is appreciated that the intrinsic virological properties of omicron strains, which exhibit enhanced replication fitness in epithelial cells, render these viruses less susceptible to neutralization *in vitro* ^58, 59, 60^. Another interesting but counterintuitive observation was that only B cells boosted by wild-type SARS-CoV-2, and not by Omicron variants, could neutralize omicron variants (Fig. 5A, C).

Although both germline reverted CR3022 mAb and the representative of the glCR3022H transgene bound to SARS-CoV, they exhibited no binding to SARS-CoV-2, suggesting that affinity maturation and accumulation of SHM are required for cross-reactivity (Fig. 5A). Indeed, only antibodies with more than 10 SHMs (H+L, aa) exhibited neutralizing activity against SARS-CoV-2 or VOCs (Fig. 5A and 5D). Not surprisingly, antibodies with at least 10 amino acid mutations also displayed higher binding affinity (Fig. 5A, Supplementary Fig.4, Supplementary Table 2). An exception was antibody EdOLB-7, which had the highest 30 mutations, but had low binding breadth and cannot neutralize to SARS-CoV-2 (Fig. 5A). An analysis of the binding affinity by biolayer interferometry of various antibodies, including those with marginal neutralization, suggested that the affinity threshold for SARS-CoV-2 neutralization by CR3022-like antibodies is approximately 20 nM (Fig. 5E).

We examined the role of heavy versus light chains on CR3022-like antibody affinity by expressing mAbs from shuffled heavy or light chain genes. Interestingly, we found that light chains from antibodies with greater affinities (< 5 nM) retained better binding affinity when paired with suboptimal heavy chains (e.g., CR3022, BoWLB-831), but not vice versa. This finding suggests that mutations in the light chain also play an important role in maintaining high binding affinity (Fig. 5F).

Further examination of SHM distributions across the heavy and light chains encoding the CR3022-like antibodies revealed substantial accumulation of mutations within the CDRs. In particular, CDRH1 in the knock-in *IGHV5-51* gene showed a high mutation frequency, with over 80% of heavy chains acquiring mutations T30I and S31T. These mutations represent convergent evolution of the glCR3022H gene by both humans and mice for SARS-CoV and SARS-CoV-2 binding, as they were also present in the parental human CR3022 antibody (Fig. 5G-H). These amino acid substitutions were previously shown to be important for CR3022 binding, particularly the key hydrophobic interaction between residue I30 and the CR3022 epitope ^20^. The T30I mutation was facilitated by a known hotspot of mutator enzyme Activation-Induced Deaminase in the germline sequence, targeting the second nucleotide of the isoleucine codon, leading to a transition from ACC (T) to ATC (I). Mutations accumulating at T31 have been shown to play a key role in interacting with P384 on the SARS-CoV-2 RBD ^20^. Similar convergent amino acid substitutions were also selected on the light chains, despite the difference between human *IGKV4-1* gene in the parental CR3022 and the murine *Igkv8-30* gene selectively used in CR3022-like antibodies in mice. While murine *Igkv8-30* differs from the human *IGKV4-1* by 14 amino acid residues, it is the closest homologous. Most of these amino acid differences are conservative substitutions that are primarily located in the framework regions (e.g., V15L, K18R, V19A). The N28I mutation in CDRL1 was observed in approximately 50% of adapted *Igkv8-30* and is also present in the parental CR3022 light chain (Fig. 5G-H). Previous structural analysis of CR3022 binding to SARS-CoV RBD showed that the N28I exchange, along with other mutations at Y27d, Y32, and W50, collectively formed a hydrophobic pocket that interacts with the side chain of M430 on the SARS-CoV RBD^20^. Notably, the M430 residue is replaced by threonine in SARS-CoV-2, and the I28 exchange often occurs after priming immunization with SARS-CoV, making it less likely to be selected during SARS-CoV-2 boosting. Over 80.4% (267/332) of the mouse-adapted heavy chain transgenes shared at least two convergent mutations with parental CR3022 (Fig. 5I). Analysis of a random reference set of 378 full-length human *IGHV5-51* cDNA sequences (Supplementary Table 3), unrelated to SARS-CoV-2 responses, found that 25.9% (98/378) of the sequences incorporated at least two amino acid mutations like those in CR3022 (Supplementary Fig.5D). These finding support convergent evolution, with selective accumulation of mutations similar to CR3022 during affinity maturation in the mouse model ^61^.

In summary, our characterization of CR3022-like antibody variants from sequentially immunized transgenic mice reveals a clear affinity maturation process involving convergent heavy and light chain mutations. Notably, additional mutations or light chain variations were also selected to enhance the binding and neutralizing potency of mouse CR30222-like antibodies against SARS-CoV-2 and related VOCs, highlighting their broad-spectrum neutralization potential. To decipher the basis of this adaptation, we performed structure-function studies on selected Fab/SARS-CoV-2 RBD complexes.

### Structural studies of three adapted CR3022-like antibodies in complex with SARS-CoV-2 RBD

To investigate the mechanism of the increased SARS-CoV-2 neutralizing activity of the *in vivo* selected antibodies at the molecular level, we determined crystal structures of three antibodies BoWLB-105, BoWLB-622, and BoWLB-1173 Fabs in complex with SARS-CoV-2 WT RBD (n.b. Fab CC12.3 was also added to the complex to aid in crystallization) at 2.54, 2.62, and 2.81 Å resolution, respectively (Fig. 6, Supplementary Table 4). We compared these three complex structures with CR3022 in complex with SARS-CoV-2 WT RBD (PDB 6W41) and eCR3022.20, which was evolved and selected by yeast display, in complex with SARS-CoV-2 WT RBD (PDB 8GF2) ^20, 27^.

**Figure. 6.**
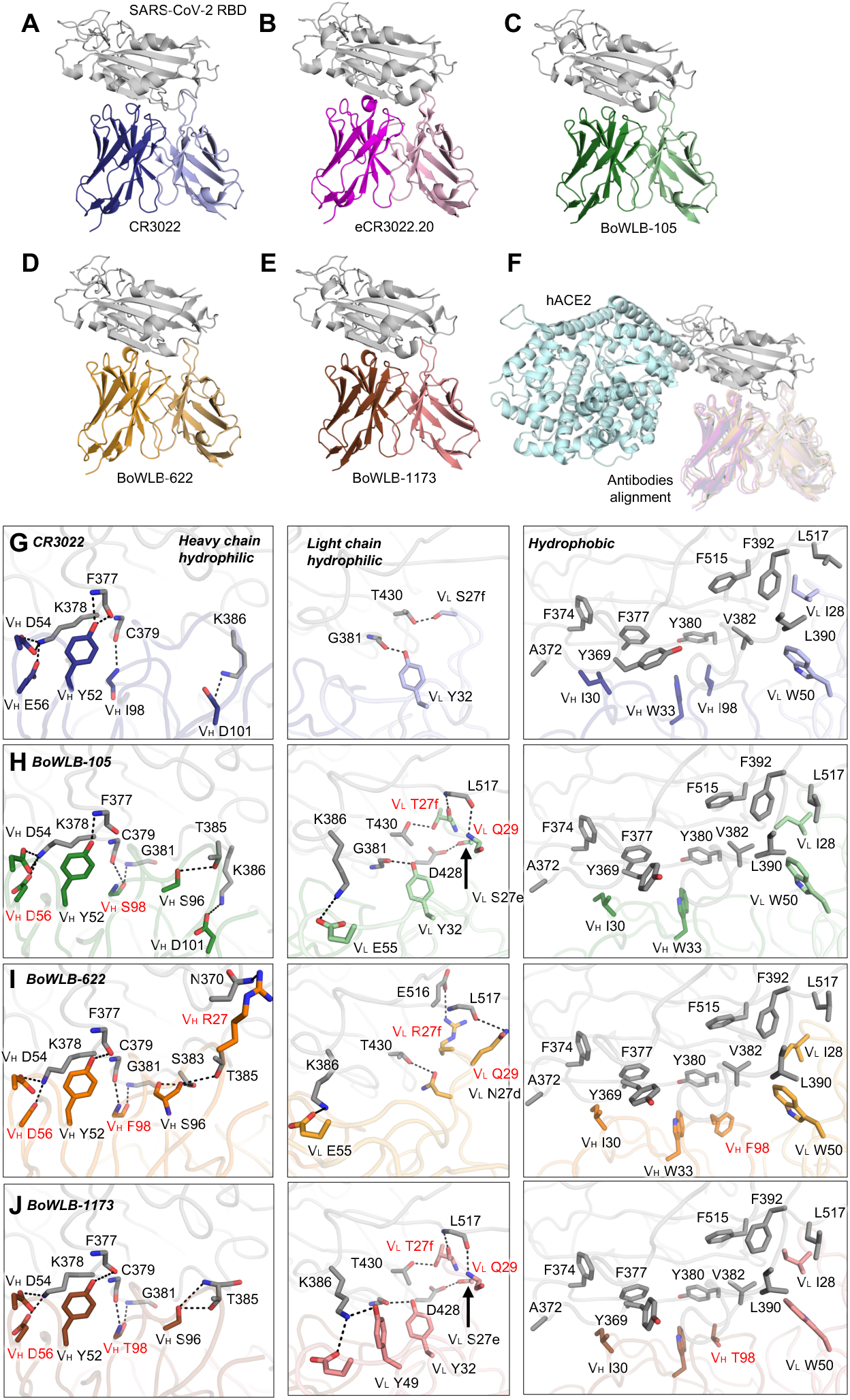
Crystal structures and detailed atomic interactions of three engineered CR3022 antibodies in complex with SARS-CoV-2 WT RBD. For clarity, only the Fab variable domains are shown. Heavy and light chains of CR3022, eCR3022.20, BoWLB-105, BoWLB-622, and BoWLB-1173 are colored in blue/lavender, magenta/pink, green/light green, orange/light orange, and brown/salmon, respectively. SARS-CoV-2 WT RBD and ACE2 is colored in grey and pale cyan, respectively. Kabat numbering is applied to the antibodies. Hydrogen bonds and salt bridges are represented by black dashed lines. Paratope residues of engineered antibodies that differ from CR3022 are highlighted in red. (A-B) Our previous crystal structures of SARS-CoV-2 RBD in complex with CR3022 andeCR3022.20 are shown for comparison (PDB: 6W41, 8GF2). (C-E) Overall view of the BoWLB-105/SARS-CoV-2 RBD, BoWLB-622/SARS-CoV-2 RBD, BoWLB-1173/SARS-CoV-2 RBD structures at 2.54 Å, 2.62 Å, and 2.81 Å, respectively. (F) Structural alignment of all antibodies using RBD as reference. ACE2-RBD complex (PDB: 6M0J) is displayed to demonstrate that there is no clash between ACE2 and these antibodies. (G-J) Detailed hydrophilic interactions (H-bonds and salt bridges, left) and hydrophobic interactions (right) of SARS-CoV-2 RBD with CR3022, BoWLB-105, BoWLB-622, and BoWLB-1173, respectively.

These three *in vivo* adapted CR3022 antibodies bind to the same epitope of SARS-CoV-2 RBD with nearly identical angles and disposition as CR3022 and eCR3022.20 (Fig. 6A-E), and do not interfere with binding to the cell receptor ACE2 (Fig. 6F). Furthermore, the RBD conformations bound by all antibodies are almost identical: Cα RMSD values were all 0.7 Å for the RBD with BoWLB-105, BoWLB-622, and BoWLB-1173, compared to SARS-CoV-2 RBD in complex with CR3022 (Supplementary Fig.6A, B, E). Among these, the RBDs in the structures complexed with BoWLB-105 and BoWLB-1173 exhibit some deviations in loop 517-520 (Supplementary Fig.6). Through structural and sequence comparisons, we found that residue 27f is serine (S) and 29 is asparagine (N) on the light chain of CR3022, while 27f on the light chains of BoWLB-105 and BoWLB-1173 is substituted by a threonine (T) and 29 by a glutamine (Q) (Supplementary Fig.6C, D, F). The side chain of glutamine is longer than that of asparagine and now forms an internal H-bond with the main-chain carbonyl of tyrosine 27d. This interaction rearranges the local conformation of CDRL1. It facilitates interactions between the main-chain amide and carbonyl of RBD L517 with the main-chain carbonyl of V_L_ T27f and amide of V_L_ Q29 of Fabs BoWLB-105 and BoWLB-1173, causing a shift in loop 517-520 as well as a change in conformation at the tip of CDRL1 (Supplementary Fig.6C and F). In contrast, this distance is too long in CR3022 to form hydrogen bonds with V_L_ S27f and V_L_ N29 (Supplementary Fig.6D).

Although the RBD is polymorphic due to the evolution of SARS-CoV-2, residues at the CR3022 epitope are relatively well conserved ^62^ (Supplementary Fig.6G). These conserved residues contribute extensively to the recognition by all three mouse-adapted CR3022-like antibodies (Fig. 6G-J), which explains their broad reactivity to many SARS-CoV-2 VOCs. For all three CR3022-like antibodies and CR3022, hydrophobic paratope residues in both the heavy and light chains interact with SARS-CoV-2 RBD conserved aromatic and aliphatic residues Y369, A372, F374, F377, Y380, V382, L390, F392, F515, and L517 (Fig. 6H-J). All antibodies contain four conserved hydrophobic paratope residues (V_H_ I30, W33, and V_L_ I28, W50) that participate in hydrophobic interactions, while V_H_ I98 in CR3022, V_H_ F98 in BoWLB-622, and the methyl group of V_H_ T98 in BoWLB-1173 participate in hydrophobic interactions.

Paratope residues of CR3022 and the mouse-adapted CR3022-like antibodies also form extensive H-bonds and salt bridges with the SARS-CoV-2 RBD. Among them, some interactions are conserved across all antibodies, including V_H_ D54 and E56 that form salt bridges with K378, and V_H_ Y52 that forms H-bonds with the main-chain of F377. There are also some unique interactions, including H-bonds and salt bridges (Table 1 and Supplementary Fig.7). Notably, CR3022 forms only two hydrogen bonds and one salt bridge/hydrogen bond, whereas the mouse-adapted antibodies exhibit many more polar interactions with the RBD. For example, BoWLB-105 makes seven hydrogen bonds and two salt bridges with the RBD, BoWLB-622 makes seven hydrogen bonds and two salt bridges, and BoWLB-1173 makes eight hydrogen bonds and one salt bridge (Fig. 6 G-J, Table 1). In total, the three mouse-adapted CR3022 antibodies exhibit similar hydrophobic interactions to CR3022 but display more pronounced polar and electrostatic interactions, including hydrogen bonds and salt bridges, due to mutation and loop shift (Supplementary Fig.7), which can explain the increase in binding affinity and neutralizing potency against SARS-CoV-2 and VOCs.

**Table 1.** Different interactions of CR3022, BoWLB-105, BoWLB-622, and BoWLB-1173 with SARS-CoV-2 WT RBD. The four columns are the four different antibodies with the RBD. The brackets ndicate whether it is an H-bond or salt bridge. Each line is the same epitope esidue. No interaction is indicated by “X”.

| SARS-CoV-2 RBD<br>& CR3022 | SARS-CoV-2 RBD<br>& BoWLB-105 | SARS-CoV-2 RBD<br>& BoWLB-622 | SARS-CoV-2 RBD<br>& BoWLB-1173 |
| --- | --- | --- | --- |
| X | X | N370-V<sub>H</sub>R27 (H-bond) | X |
| C379-V<sub>H</sub>I98 (H-bond) | C379-V<sub>H</sub>S98 (H-bond) | C379-V<sub>H</sub>T98 (H-bond) | C379-V<sub>H</sub>F98 (H-bond) |
| X | G381-V<sub>H</sub>S98 (H-bond) | G381-V<sub>H</sub>T98 (H-bond) | G381-V<sub>H</sub>F98 (H-bond) |
| X | X | S383-V<sub>H</sub>S96 (H-bond) | X |
| X | T385-V<sub>H</sub>S96 (H-bond) | T385-V<sub>H</sub>S96 (H-bond) | T385-V<sub>H</sub>S96 (H-bond) |
| K386-V<sub>H</sub>D101 (salt bridge) | K386-V<sub>H</sub>D101 (salt bridge)<br>K386-V<sub>L</sub>E55 (salt bridge) | K386-V<sub>L</sub>E55 (salt bridge) | K386-V<sub>L</sub>E55 (salt bridge)<br>K386-V<sub>L</sub>Y49 (H-bond) |
| X | D428-V<sub>L</sub>S27e (H-bond) | X | D428-V<sub>L</sub>S27e (H-bond) |
| T430-V<sub>L</sub>S27f (H-bond) | T430-V<sub>L</sub>T27f (H-bond) | T430-V<sub>L</sub>N27d (H-bond) | T430-V<sub>L</sub>T27f (H-bond) |
| X | X | E516-V<sub>L</sub>R27f (salt bridge) | X |
| X | L517-V<sub>L</sub>Q29 (H-bond)<br>L517-V<sub>L</sub>T27f (H-bond) | L517-V<sub>L</sub>Q29 (H-bond) | L517-V<sub>L</sub>Q29 (H-bond)<br>L517-V<sub>L</sub>T27f (H-bond) |

## Discussion

Since the mAb CR3022 demonstrated cross-reactivity between SARS-CoV and SARS-CoV-2, it attracted significant attention at the onset of the COVID-19 pandemic as one of the early examples of broadly-conserved epitopes on coronavirus spikes ^19^. However, a key mutation P384A in the SARS-CoV-2 RBD rendered CR3022 ineffective at neutralizing the virus, although it still maintained some binding affinity ^23^. Attempts from other groups to explore the possibilities of affinity enhancement of CR3022 to SARS-CoV-2 RBD using *in vitro* mutagenesis or computationally have been reported ^27, 28, 29, 30, 31, 32^. While this manuscript was in preparation, *Nair et al*. independently demonstrated that *in vivo* affinity maturation can enable CR3022 to acquire SARS-CoV-2 neutralizing activity ^63^. However, it is not clear how easily the CR3022 epitope can be targeted *in vivo*, and despite reports of thousands of mAbs to SARS-CoV-2 spike from patients, examples of such antibodies are rare ^64, 65^. In this study, we deployed Ig-humanized knock-in mice to experimentally assess whether predicted germline CR3022-like B cell precursors at physiological frequency can be efficiently adapted across betacoronavirus strains. We reported that CR3022-like B cell precursors can be activated and recruited to GCs when congenic recipient mice are primed with the SARS-CoV spike protein, but not with the SARS-CoV-2 spike. These results suggest that the induction of CR3022-like antibodies may require prior exposure to spike components from other SARS-related viruses that meet the necessary affinity threshold for priming. This observation may account for the rarity of detected human mAbs to the CR3022 epitope in past studies. Thus, developing a multivalent or mosaic coronavirus vaccine could target not only SARS-related viruses responsible for past epidemics and pandemics but also those with the potential to cause future global health crises ^66, 67^.

We have developed the glCR3022H mouse model to study the targeting of an important conserved class of coronavirus spike epitope, with the additional goal of integrating it into a valuable experimental platform for immunology studies. Most mouse-model systems for studying immunoglobulin affinity maturation and germinal center biology involve the use of simplified and highly predictable hapten-specific responses, such as to 4-hydroxy-3-nitrophenylacetyl (NP) ^68^, with contrived T cell help provided by coupling the haptens to carrier proteins such as OVA. While these systems allow for facile measurement of affinity maturation, they lack the complex binding interfaces typically seen between antibodies and proteinaceous antigens and accordingly cannot model the critical, and often subtle, chain of mutational events leading to the production of high-quality antibody. Furthermore, hapten systems lack competitive interactions with other epitopes and are not directly applicable to pathogen immunity. Herein, we report a powerful model for studying B cell responses and affinity maturation in a highly predictable system with simple experimental readouts and against complex and relevant protein antigens. By comparing SARS-CoV or SARS-CoV-2 spike binding with or without a competitive CR3022 Fab, binding to the CR3022 epitope is easily measured in simple flow cytometry or linking B cell receptor to antigen specificity by sequencing (LIBRA-seq) ^34^ assays. Using these probes after SARS-CoV spike priming and SARS-CoV-2 booster immunization, one can readily detect frequencies of B cells with successful affinity maturation, as only glCR3022H genes with accumulated somatic mutations and selection can bind to the CR3022 epitope on SARS-CoV-2 spikes. Furthermore, only B cells with the glCR3022H paired with *Igkv8-30* bind to the CR3022 epitope, enabling a rapid prediction tool for epitope specificity through simple sequencing. This also simplifies the selection of cells for mAb production, or single-cell transcriptome analysis, for example. Finally, the convergent evolution of amino acid substitutions that accumulate in CR3022-like B cells in these mice allows a highly granular or precise means to evaluate successful affinity maturation. Thus, this mouse model offers a powerful and quantifiable system for studies on GC responses and affinity maturation aimed at testing the role of various B cell-expressed proteins (knockouts or transgenes), the role of other effector cells or molecules, or even manipulations affecting gross histological organization.

Using these mice, we observed that B cell precursor frequency does not correspond to immunodominance, as neither a high number of transgenic glCR3022H B cells before immunization nor early (day 7) induction of abundant GC B cells targeting epitopes on the spike overlapping with CR3022 in non-transgenic mice, which account for 30% of the cells, ultimately led to the dominant targeting of the CR3022 epitope. Unexpectedly, we observed that immune responses to the immune-subdominant CR3022 epitope plateaued 3-4 weeks in the transgenic mice or plummeted in non-transgenic mice after the first boost with the SARS-CoV-2 spike protein. Despite additional boosts, the immune response appeared to shift toward a diverse set of epitopes within the RBD, as evidenced by serological and Ig repertoire results. This effect was particularly pronounced with escalating dosages, consistent with previous findings that escalating dosages enhance GC responses and modulate the immunodominance hierarchy ^35, 50^. Indeed, compared to conventional bolus immunizations, we found that escalating dosage immunization required less time to achieve a saturated serum IgG titer against the SARS-CoV-2 RBD. We suspect that antibody feedback or potential epitope masking may play a significant role in limiting the CR3022 epitope-specific B cell response, leading to a more diverse immune response targeting other, more immunodominant epitopes on the RBD, such as those surrounding the RBS ^69, 70^. Thus, vaccine strategies aimed at targeting particular immuno-subdominant epitopes, which are often the most conserved epitopes on mutating viral antigens, may need to focus on effectively maintaining epitope-specific GC reactions despite existing serum antibodies against the epitopes. This could be accomplished by iterative boosts using immunogens with similar but not identical presentations of an epitope, similar to germline targeting strategies in the HIV field ^71^, allowing continued affinity maturation.

Previous studies on engineering CR3022 mAbs primarily employed site-directed mutagenesis *in vitro*, with mutations largely confined to the CDRs of CR3022 ^27, 30^. In this study, by utilizing SHM targeting *in vivo*, we demonstrated convergent evolution of key mutations in amino acid residues of CR3022, particularly those involved in major hydrophobic interactions with its epitope, were well-replicated in CR3022-like clones. Among the conserved mutations, I30 and T31 in CDRH1 appeared to be under the greatest positive selective pressure. However, we and others also found that CR3022-like mAbs that lack these two key mutations may still retain neutralizing activity, as exemplified by mAb EdOLB-1, which harbors N30 and L31 instead, or CR3022 antibodies engineered *in vitro* that contain W31 ^27^. We noted that an affinity threshold for neutralization of SARS-CoV-2 at the CR3022 epitope is approximately 20 nM, while the upper limit of affinity *in vivo* of CR3022-like clones approached 1 nM. While selection for conserved residues was typically evident, mouse-adapted CR3022 antibodies had readily improved affinity over this threshold to neutralize SARS-CoV-2 and VOCs through various mutations that resulted in the formation of additional hydrogen bonds and salt bridges.

In conclusion, our results demonstrate that under a sequential SARS-CoV/SARS-CoV-2 prime-boost immunization regimen, CR3022-like B cells can be effectively activated, participate in the germinal center response, and accumulate on-track SHMs to achieve broad neutralization. Moreover, a conventional bolus dosing strategy for coronavirus vaccination may be preferable to prevent the excessive diversion of humoral responses away from certain subdominant yet protective epitopes.

## Materials and Methods

### Ethics statement

All animal experiments were conducted in compliance with protocols approved by the Institutional Animal Care and Use Committee (IACUC) of Weill Cornell Medicine (Protocol #2021-0024) and adhered to the guidelines of the Association for Assessment and Accreditation of Laboratory Animal Care International (AAALAC). Both male and female mice aged 6–12 weeks were used and maintained under specific pathogen-free conditions.

### Generation of glCR3022H knock-in mouse

glCR3022H knock-in mice were generated as previously described ^36^. Briefly, the Speed-Ig donor DNA construct was assembled by combining SalI-linearized intermediate constructs, the promoter region, and the germline-reverted CR3022 heavy chain variable region. Assembly was performed using NEB HiFi 2× Master Mix according to the manufacturer’s instructions. Following confirmation by Sanger sequencing, the donor DNA was diluted in ultrapure water to a final volume of 100 µL, containing 50 ng/µL crRNA:tracrRNA, 50 ng/µL Cas9 protein (IDT), and 15 ng/µL targeting construct DNA. Fertilized mouse zygotes were microinjected with the prepared mix and implanted into pseudopregnant females. F₀ pups were genotyped to identify knock-in founders. Positive F₀ glCR3022H mice were crossed with WT C57BL/6J females to produce F₁ progeny and establish the colony. Genotyping was performed on ear or tail biopsies using conventional PCR.

### Adoptive transfer

B6.SJL-*Ptprc*^a^*pepc*^b^/BoyJ (CD45.1^+^) recipient mice aged 8-12 weeks were purchased from Jackson Laboratory. Donor B cells (CD45.2^+^) were isolated from male or female glCR3022H donor knock-in mice aged 8-16 weeks using a mouse Pan-B cell negative selection kit (StemCell Technologies). Purified cells were resuspended in sterile PBS at the desired concentration and then adoptively transferred into same-sex CD45.1^+^ recipient mice via retro-orbital injection.

### Immunizations

Spike proteins from either SARS-CoV or SARS-CoV-2 were administered intraperitoneally at a dose of 10 μg per mouse in a bolus immunization regimen, diluted in PBS to a final volume of 100 μL. For the escalating-dose regimen, the same total antigen amount (10 μg) was divided across seven injections given over a two-week period, with doses administered every other day as previously described ^35^. The individual doses from the 1st to 7th immunization were 0.02, 0.04, 0.12, 0.32, 0.86, 2.33, and 6.33 μg, respectively. Immunogen solutions were mixed 1:1 with Sigma Adjuvant System (Ribi) and incubated for 30 minutes before injection. The final formulation (200 μL total volume) was administered intraperitoneally.

### Recombinant antigen generation

All recombinant antigens were produced in Expi293F cells (Thermo Fisher Scientific) as previously described ^72^. Briefly, plasmids encoding full-length spike or receptor-binding domain (RBD) proteins from zoonotic strains, SARS-CoV, SARS-CoV-2 and relevant VOCs were transiently transfected into Expi293F cells using a standard transfection protocol. Secreted proteins were harvested from the supernatant and purified via Ni-NTA affinity chromatography (Qiagen). Purified proteins were subsequently buffer exchanged into PBS using centrifugal filtration.

### Flow cytometry and probe generation

For B cell compartment analysis, bone marrow cells were harvested by flushing femurs and tibiae with sterile DPBS, followed by red blood cell lysis using ACK lysing buffer (Quality Biological). Spleens were mechanically dissociated through 70-μm cell strainers (BD Biosciences) into single-cell suspensions, and erythrocytes were lysed using ACK buffer. Peritoneal cavity cells were collected by lavage, taking care to avoid blood contamination. All cells were resuspended in FACS buffer (PBS supplemented with 2% FBS) and incubated with Fc-blocking antibody (clone 2.4G2, Bio X Cell). Different fluorescent antibody panels were used for bone marrow cells [CD19-PE, B220-FITC, IgM-PE Cy7, IgD-BUV395, CD43-AF700, CD24-APC, CD249(BP-1)-BV786, CD93-BV711, CD3-PB, CD11b-PB, Gr-1-PB, F4/80-PB], splenocytes (CD19-PE, CD43-AF700, CD23-APC, CD21/35-APC Cy7) and peritoneal cells (IgM-PE Cy7, CD11b-PB, CD19-PE, CD23-APC, CD5-BV711) at 1/100 or 1/200 dilutions. 7-Aminoactinomycin D (7-AAD) was added to each sample for viability staining. Flow cytometric acquisition was performed on a BD LSRFortessa (BD Biosciences) or a Cytek Aurora (Cytek Biosciences), and data were analyzed using FlowJo software (BD).

For characterization and sorting of antigen-specific B cells, Avi-tagged SARS-CoV-2 RBD or hemagglutinin (HA, control) proteins were expressed in-house, biotinylated using a BirA-500 kit (Avidity), and tetramerized by conjugation with fluorophore-labeled streptavidin (BioLegend) at a 4:1 molar ratio. Full-length spike protein was biotinylated using EZ-Link NHS-PEG4-Biotin (Thermo Scientific). Cells were stained with antigen probes and fluorophore-conjugated antibodies in FACS buffer supplemented with 2 mM free biotin. Antigen-specific B cells were sorted into 96-well plates or 1.5 mL microcentrifuge tubes using a BD FACSMelody cell sorter or Cytek Aurora CS system for downstream applications.

### Single cell PCR and Sanger sequencing

Following single-cell sorting of antigen-specific B cells, variable region genes of the heavy and light chains were amplified by reverse transcription PCR. First-strand cDNA synthesis was carried out using SuperScript IV First-Strand Synthesis System (Invitrogen) following the manufacturer’s instructions. Nested PCR, consisting of round 1 and round 2 reactions, was performed in 25-μL volumes using 2X DreamTaq Master Mix (Thermo Fisher), along with Ig heavy– and kappa chain– specific primer mixes under thermocycling conditions previously described ^42, 73^. PCR products were resolved on 1.5% agarose gels stained with GreenGlo DNA dye (NEB), and wells with correctly sized bands were submitted to GENEWIZ for Sanger sequencing. Sequence analysis and alignment were performed using IMGT/V-QUEST (http://www.imgt.org).

### scRNA-seq library preparation and data analysis

Sorted B cells were assessed for viability using Trypan blue and immediately processed for 10x Genomics GEM generation using a Chromium X Controller. Mouse B cell V(D)J, 5’ gene expression, and feature barcode libraries were prepared according to the manufacturer’s protocol. Libraries were pooled and sequenced on an Illumina NextSeq 1000 instrument using parameters recommended by 10x Genomics. Demultiplexing was performed using the Cell Ranger mkfastq pipeline (v7.0.0–v9.0.0), and FASTQ files were aligned to the mouse reference genome (mm10-2020-A) using the Cell Ranger count and VDJ pipelines. The reference for VDJ analysis was customized to include exogenous knock-in VDJ genes.

Downstream gene expression analysis was carried out in R (v4.4.3) using Seurat (v4.3.0 or newer). Cells were filtered based on gene count (retaining cells with 200–2,500 genes) and percentage of mitochondrial gene expression. Full-length V(D)J contigs were assembled using Cell Ranger VDJ or Cell Ranger multi (v7.0.1 or newer).

### Monoclonal antibody production and binding measurements by bio-layer interferometry

Monoclonal antibody (mAb) heavy and light chain genes were synthesized (Twist Bioscience) and cloned into expression vectors encoding human IgG1 heavy chain, kappa light chain, or Fab fragments with C-terminal His-tags using HiFi DNA Assembly Master Mix (New England Biolabs). Expi293F cells were transiently transfected with plasmids encoding the antibody chains, and expressed mAbs and Fabs were purified using Protein A resin (GenScript) and Ni-NTA affinity chromatography (Qiagen), respectively, as previously described^73^.

Bio-layer interferometry (BLI) was performed using an Octet K2 instrument (ForteBio), and data were analyzed with Octet Analysis Studio software (Sartorius). Site-specific biotinylated receptor-binding domain (RBD) proteins were immobilized on High Precision Streptavidin (SAX) Biosensors (ForteBio) at 20 – 40 μg/mL for 5 minutes in kinetics buffer (PBS with 0.01% BSA and 0.002% Tween-20). Sensors were then transferred to wells containing serial dilutions of Fab or IgG for 120 seconds (association), followed by 120 seconds in kinetics buffer (dissociation). Apparent dissociation constants (K_D_) were determined using a global fit 1:1 binding model provided in the analysis software.

### Enzyme-linked immunosorbent assay (ELISA)

To evaluate mAb binding, recombinant antigens were coated onto high-binding 96-well polystyrene plates (Costar) at 2 μg/mL in PBS (50 μL per well) and incubated overnight at 4 °C. Plates were washed three times with PBS containing 0.05% Tween-20 and blocked with PBS supplemented with 20% FBS for 1 hour at 37 °C. Serial 3-fold dilutions of mAbs, starting at 10 μg/mL, were prepared in PBS and incubated on the antigen-coated plates for 1 hour at 37 °C. After three additional washes, HRP-conjugated goat anti-human IgG (Jackson ImmunoResearch) diluted 1:1,000 in PBS was added and incubated for detection. Plates were developed using Super Aquablue ELISA substrate (Thermo Fisher), and absorbance was measured at 405 nm using a microplate reader (Bio-Rad). To quantify antigen-specific serum IgG titers, 3-fold serial dilutions of mouse sera (starting at 1:100) were incubated at room temperature (RT) for 2 hours on plates pre-coated with antigen. Plates were pre-blocked with 3% fat-free dry milk in PBS containing 0.1% Tween-20 for 1 hour at RT prior to serum incubation. Wells were developed using o-phenylenediamine dihydrochloride (OPD; Sigma) for 10 minutes at RT, and the reaction was stopped with 3 M HCl. Absorbance was measured at 490 nm. Binding curves and area under the curve (AUC) values were calculated using GraphPad Prism 10.

### Competition ELISA

To assess antibody competition, 96-well high-binding plates were coated with 50 μL of antigen (1 μg/mL in PBS) and incubated overnight at 4 °C. After blocking with PBS containing 20% FBS for 1 h at 37 °C, mAbs (20 μg/mL) were added and incubated for 2 h at room temperature (RT). Biotinylated mAb CR3022 using EZ-Link Sulfo-NHS-Biotin (Thermo Fisher) was added at concentrations twice its Kd value and incubated for an additional 2 h at RT. Following incubation, plates were washed and developed with HRP-conjugated streptavidin (Southern Biotech) for 1 h at 37 °C. Super Aquablue substrate (Thermo Fisher) was added and absorbance was measured at 405 nm using a microplate spectrophotometer (Bio-Rad). To standardize the assays, biotinylated competitor mAb CR3022 was added to control wells without any competing antibodies, and data were recorded when the absorbance of these wells reached 1.0–1.5 OD units at 405 nm. All monoclonal antibodies were tested in duplicate, and each experiment was independently repeated. Percent competition was calculated by comparing the optical density (OD) of each sample to the corresponding control OD. To determine the AUC ratio of mouse serum (i.e., the ratio of AUC in the absence versus presence of Fab), CR3022-encoding Fab was added to antigen-coated wells at a final concentration of 5 μg/mL after the blocking step to competitively block the CR3022 epitope on the RBD. Following a 1-hour incubation at 37 °C, the Fab solution was washed off, and diluted mouse serum (starting at 1:100 in PBS) was added and incubated at room temperature (RT) for 2 hours. The subsequent ELISA color development was performed following the same procedure as the standard serum ELISA.

### Monoclonal antibody pseudo-typed virus neutralization assay

Neutralization assays were performed using SARS-CoV, SARS-CoV-2 and relevant VOCs spike-pseudotyped virus generated from recombinant replication-deficient vesicular stomatitis virus (VSV) lacking the VSV-G glycoprotein gene and encoding a luciferase reporter (Kerafast). Vero E6 cells stably expressing human ACE2 and TMPRSS2 (BEI Resources) were seeded into white, flat-bottom 96-well plates (Corning) at 40,000 cells per well and incubated overnight at 37 °C with 5% CO₂.

Monoclonal antibodies (mAbs) were four-fold serially diluted in culture medium, starting from 100 μg/mL to 0.39 μg/mL. Diluted mAbs were mixed 1:1 with pseudovirus (325–1,300 TCID₅₀/mL) and incubated at 37 °C for 1 h before addition to the Vero E6-ACE2-TMPRSS2 monolayers. Plates were then incubated for 18-24 hours at 37 °C with 5% CO₂. Following infection, supernatants were removed, and cells were lysed using luciferase assay reagent (Promega). Luminescence was measured using a SpectraMax M5 microplate reader (Molecular Devices). Half-maximal inhibitory concentrations (IC₅₀) were calculated using a custom Microsoft Excel macro.

### Focus reduction neutralization test (FRNT)

A 2-fold dilution series of antibodies starting at a final concentration of 400 µg/mL was mixed with approximately 800 focus-forming units of the virus and incubated for 1 h at 37 °C. The antibody-virus mixture was inoculated onto Vero E6/TMPRSS2 cells in 96-well plates and incubated for 1 h at 37 °C. An equal volume of 1% methylcellulose solution was added to each well, and then the cells were incubated for 16 h at 37 °C and then fixed with formalin. Foci that formed were immunostained with a mouse monoclonal antibody against SARS-CoV-2 nucleoprotein (clone 1C7C7, Sigma-Aldrich, catalog #MA5-29982, 1:10,000 dilution), followed by a horseradish peroxidase-labeled goat anti-mouse immunoglobulin (ThermoFisher, catalog #31430, 1:2500 dilution) and then were stained with TrueBlue Substrate (SeraCare Life Sciences). After plates were dried, the number of foci were quantified by using an ImmunoSpot S6 Analyzer, ImmunoCapture software, and BioSpot software (Cellular Technology).

### Crystal structure determination

The receptor-binding domain (RBD) (residues 334-528) of the SARS-CoV-2 spike (S) protein (GenBank: QHD43416.1) was cloned into a customized pFastBac vector. The RBD constructs were fused with an N-terminal gp67 signal peptide and a C-terminal His_6_ tag. Recombinant bacmid DNA was generated using the Bac-to-Bac system (Life Technologies). Baculovirus was generated by transfecting purified bacmid DNA into Sf9 cells using FuGENE HD (Promega), and subsequently used to infect suspension cultures of High Five cells (Life Technologies) at an MOI of 5 to 10. Infected High Five cells were incubated at 28 °C with shaking at 110 r.p.m. for 72 h for protein expression. The supernatant was then concentrated using a 10 kDa MW cutoff Centramate cassette (Pall Corporation). The RBD protein was purified by Ni-NTA, followed by size exclusion chromatography, and buffer exchanged into 20 mM Tris-HCl pH 7.4 and 150 mM NaCl.

Purified BoWLB-105/622/1173 Fab, CC12.3 Fab, and SARS-CoV-2 RBD were mixed at an equimolar ratio and incubated overnight at 4°C. The complex (12 mg/ml) was screened for crystallization with the 384 conditions of the JCSG Core Suite (QIAGEN) on our robotic CrystalMation system (Rigaku) at Scripps Research by the vapor diffusion method in sitting drops containing 0.1 μl of protein and 0.1 μl of reservoir solution. Diffraction-quality crystals were obtained in the following conditions: BoWLB-105-SARS-CoV-2 RBD-CC12.3 and BoWLB-1173-SARS-CoV-2 RBD-CC12.3: 0.1 M sodium citrate–citric acid buffer (pH 4.0), 25% (v/v) polyethylene glycol 200, and 5% (w/v) polyethylene glycol 6000 at 20°C. BoWLB-622-SARS-CoV-2 RBD-CC12.3: 0.1 M sodium citrate–citric acid buffer (pH 5.0), 1.6 M ammonium sulfate, and 20% (v/v) glycerol at 20°C.

Crystals appeared on day 3 and were harvested on day 10. The crystals were then flash-cooled and stored in liquid nitrogen until data collection. No additional cryoprotectant was required. Diffraction data were collected at cryogenic temperature (100 K) at National Synchrotron Light Source II (NSLS-II) beamlines 17-ID-1 and 17-ID-2 with beam wavelengths of 0.92009 Å and 0.97934 Å, respectively. Diffraction data were processed with HKL2000 ^74^. Structures were solved by molecular replacement using PHASER ^75^. Iterative model building and refinement were carried out in COOT ^76^ and PHENIX ^77^, respectively. Epitope and paratope residues, as well as their interactions, were identified by accessing PISA at the European Bioinformatics Institute (http://www.ebi.ac.uk/pdbe/prot_int/pistart.html) ^78^.

### Quantification and statistical analyses

All statistical analyses were performed using Prism 10 (GraphPad). Sample sizes and statistical tests are indicated in the corresponding figure legends. Data were considered statistically significant at P < 0.05.

## Acknowledgments

We thank Thomas Miller from the Gale and Ira Drukier Institute for Children’s Health at Weill Cornell Medicine for help with cell sorting. We thank the Colony Management Group from the Zuckerman Research Center and the animal technicians at the Belfer Research Building for managing the mouse colonies. We thank Florian Krammer from the Icahn School of Medicine at Mount Sinai for providing the recombinant antigens of HKU3-8 and BM48-31 strains. We thank Erik Procko from the University of Illinois at Urbana-Champaign for providing us the plasmid encoding recombinant RBD of Rs4231 strain. The Wilson lab is grateful to the staff of the National Synchrotron Light Source II (NSLS-II) beamlines 17-ID-1 and 17-ID-2 for assistance. This research used resources of NSLS-II, a U.S. Department of Energy (DOE) Office of Science User Facility operated for the DOE Office of Science by Brookhaven National Laboratory under Contract No. DE-SC0012704. We thank Henry Tien for technical support with the crystallization robot. We thank Robyn Stanfield for assistance in data collection and Marc-André Elsliger with computation.

This study was funded in part by the National Institute of Allergy and Infectious Diseases (NIAID), National Institutes of Health grant numbers U19AI082724 (PCW), U19AI109946 (PCW), U19AI057266 (PCW), R01AI190286 (MY and IAW), the NIAID Centers of Excellence for Influenza Research and Surveillance (CEIRS) grant number HHSN272201400005C (PCW), the NIAD Centers of Excellence for Influenza Research and Response (CEIRR) grant number 75N93019R00028 (PCW), and the Gates Foundation INV-004923 (IAW). This work was also partially supported by the NIAID Collaborative Influenza Vaccine Innovation Centers (CIVIC; 75N93019C00051, PCW). Y.K. and P.C.W. were funded by NIAID’s Pan-Coronavirus Vaccine Development Program (P01AI165077).

## Author contributions

Y.F., Z.F., S.E., I.A.W. and P.C.W. conceptualized the study. Y.F., Z.F., S.A.E., L.L., J.C.C., C.A.T., J.S., A.Y. and S.C. developed methodology. Y.F., Z.F., S.A.E., P.J.H., L.L., J.C.C., C.A.T., J.S., A.Y., S.C., M.H. and N.-Y.Z. conducted investigation. Y.F., Z.F., L.L., J.C.C. and C.A.T. performed visualization. M.Y., Y.K., I.A.W. and P.C.W. acquired funding. I.A.W. and P.C.W. supervised the study. Y.F., Z.F. and P.C.W. prepared the first draft. All authors reviewed and edited the manuscript.

## Competing interests

The authors declare no competing financial interests.

**Supplementary Figure 1.**
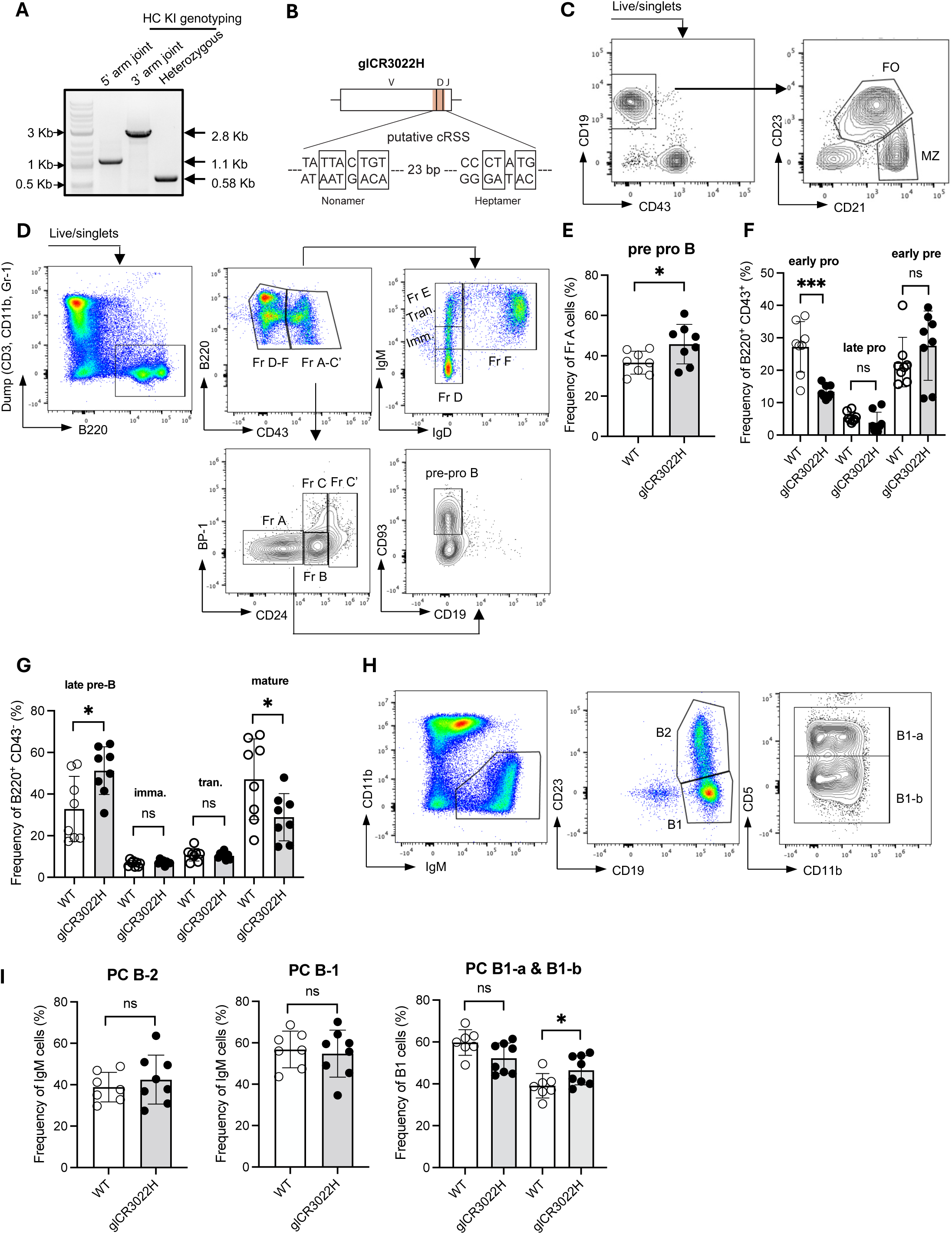
Characterization of B cell compartments in bone marrow, spleen and peritoneal cavity in gCR3022H knock-in mouse model. **(A)** Genotyping of glCR3022H knock-in founders using the 5’ and 3’ primers depicted in Figure 1A reveals a 1.1 kb and 2.8 kb amplicon spanning 5’ and 3’ homology arm joint, respectively. A 580 bp band in the third lane(heterozygous) suggests the endogenous IgH allele relative to the targeted allele remains intact, which indicates the founder is heterozygous. All primer sequences are listed in Table S1. **(B)** The location of cRSS (in pink color) was found within CDRH3 of KI sequence. The boxes enclose the bases that composed consensus sequences conserved across the canonical 23-RSSs; **(C-D)** Representative flow cytometry plots showing the gating strategy for follicular and marginal zone B cell subsets in spleen, early B cell subsets in bone marrow. The nomenclature for each fraction of B cell populations is: Fr A-C’ (B220^+^ CD43^+^); Fr A [pre pro-B (CD19^-^ CD93^+^)]; Fr B [early pro-B (CD24^int^ BP-1^-^)]; Fr C [late pro-B (CD24^int^ BP-1^+^)]; Fr C’ [early pre-B (CD24^+^ BP-1^+/-^)]; Fr D-F (B220^+^ CD43^dim/neg^); Fr D [late pre-B (IgM^-/low^ IgD^-^)]; Fr E [imm. & tran. B (IgM^int/high^ IgD^-^)]; Fr F [recirculating mature B (IgM^int/high^ IgD^+^)]. **(E-G)** Quantification of B cell progenitors in bone marrow as shown in (C). **(H)** Flow cytometry plots for characterizing B cell population in the peritoneum. **(I)** Quantification of B cell subsets as shown in (G). Error bars in all quantification plots indicate mean ± SD, and data were pooled from two independent experiments. \**p*<0.05, \*\**p*<0.01, \*\*\**p*<0.001(two-tailed Student’s *t* test). Each dot represents one mouse.

**Supplementary Figure 2.**
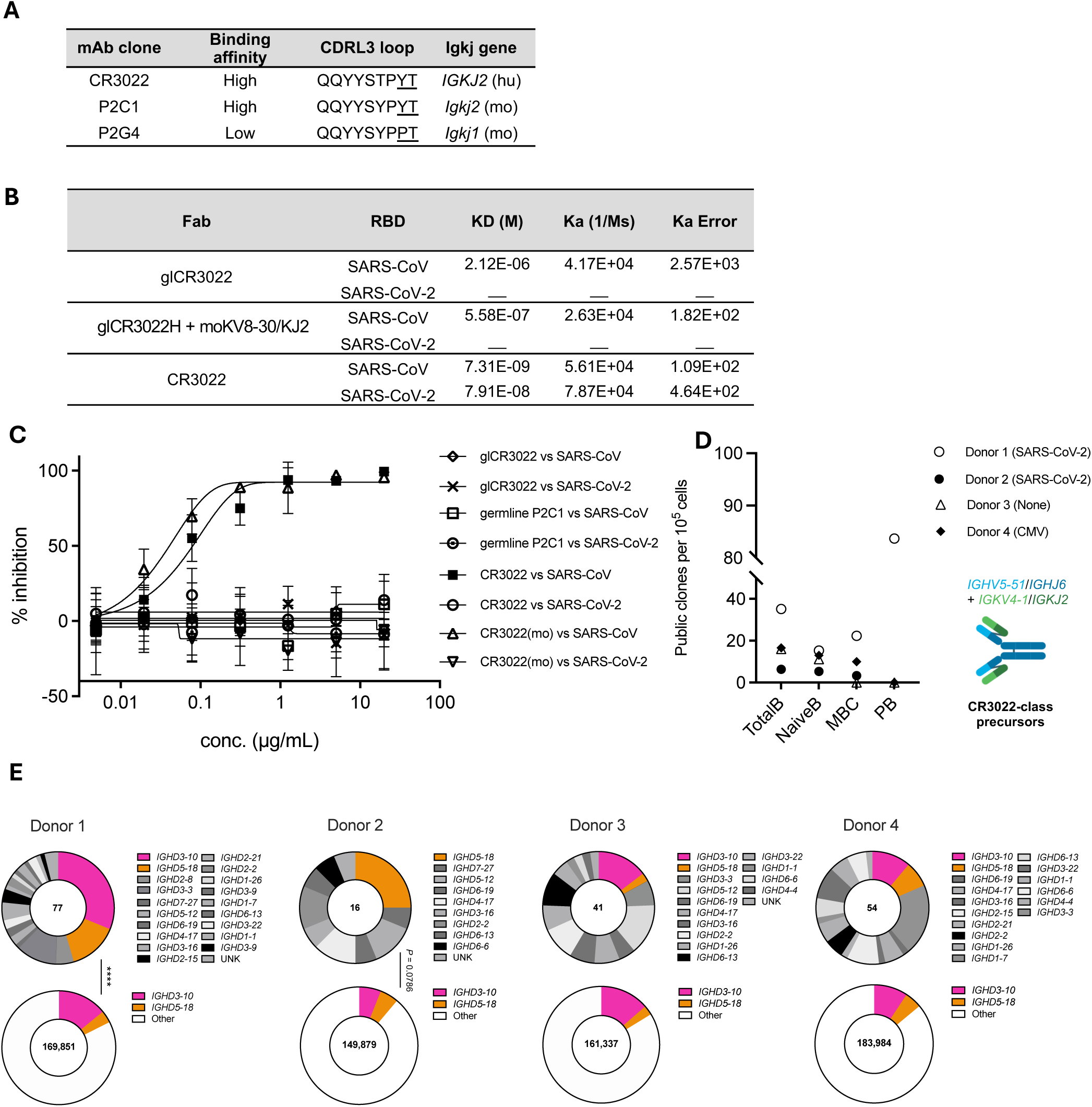
Monomeric affinities of germline and affinity-matured CR3022 and frequency of CR3022 precursors among human population. **(A)** The CDRL3 amino acid sequences and *Igkj* genes used by the three mAb clones; **(B)** The affinities of Fab of germline CR3022, germline CR3022 heavy chain paired with mouse *Igkv8-30* and Fab of affinity-matured CR3022 to SARS-CoV and SARS-CoV-2 RBDs measured by biolayer interferometry; **(C)** Neutralization curves of CR3022 and transgene-encoded CR3022 [denoted by (mo)] both in germline and affinity-matured forms tested with pseudo-typed SARS-CoV and SAR-CoV-2 viruses. **(D)** Frequencies of the CR3022-calss antibody precursors among four B cell subsets of 4 human donors. The criteria for defining CR3022-class precursors is that the BCRs are composed of heavy chain using *IGHV5-51*/*IGHJ6* and light chain using *IGKV4-1*/*IGKJ2*; **(E)** Pie charts of proportions of *IGHD3-10* and *IGHD5-18* usage among naïve B cell populations (bottom panels) and CR3022-class clones (top panels) from all four B cell subsets of the 4 human donors shown in panel **D**. The numbers of naïve B cells and CR3022-class clones analyzed are indicated in the centers of the charts. \*\*\*\**p* < 0.0001 between the D gene usage proportions for donor 1 analyzed by Chi-square test.

**Supplementary Figure 3.**
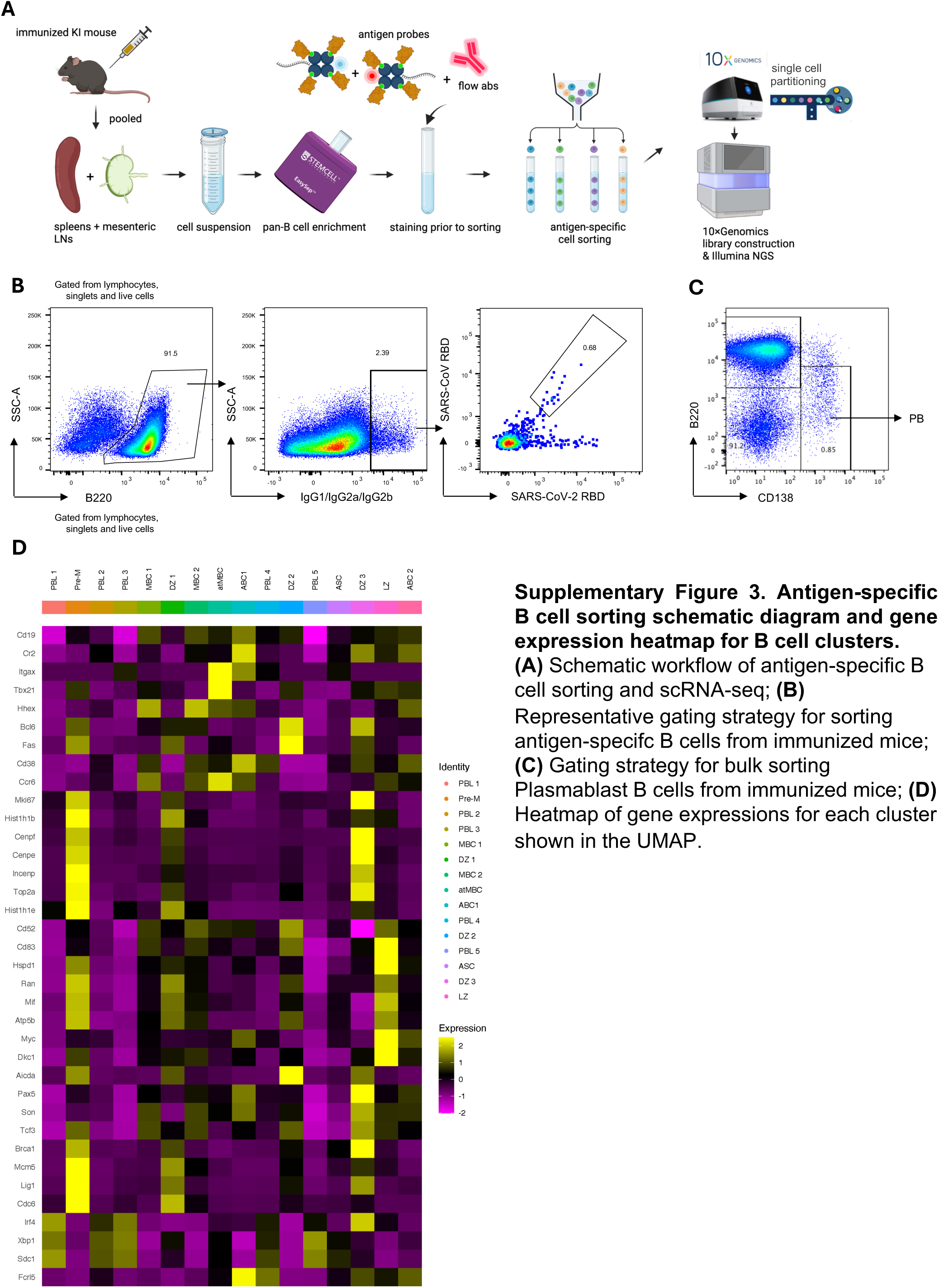
Antigen-specific B cell sorting schematic diagram and gene expression heatmap for B cell clusters. **(A)** Schematic workflow of antigen-specific B cell sorting and scRNA-seq; **(B)** Representative gating strategy for sorting antigen-specifc B cells from immunized mice; **(C)** Gating strategy for bulk sorting Plasmablast B cells from immunized mice; **(D)** Heatmap of gene expressions for each cluster shown in the UMAP.

**Supplementary Figure 4.**
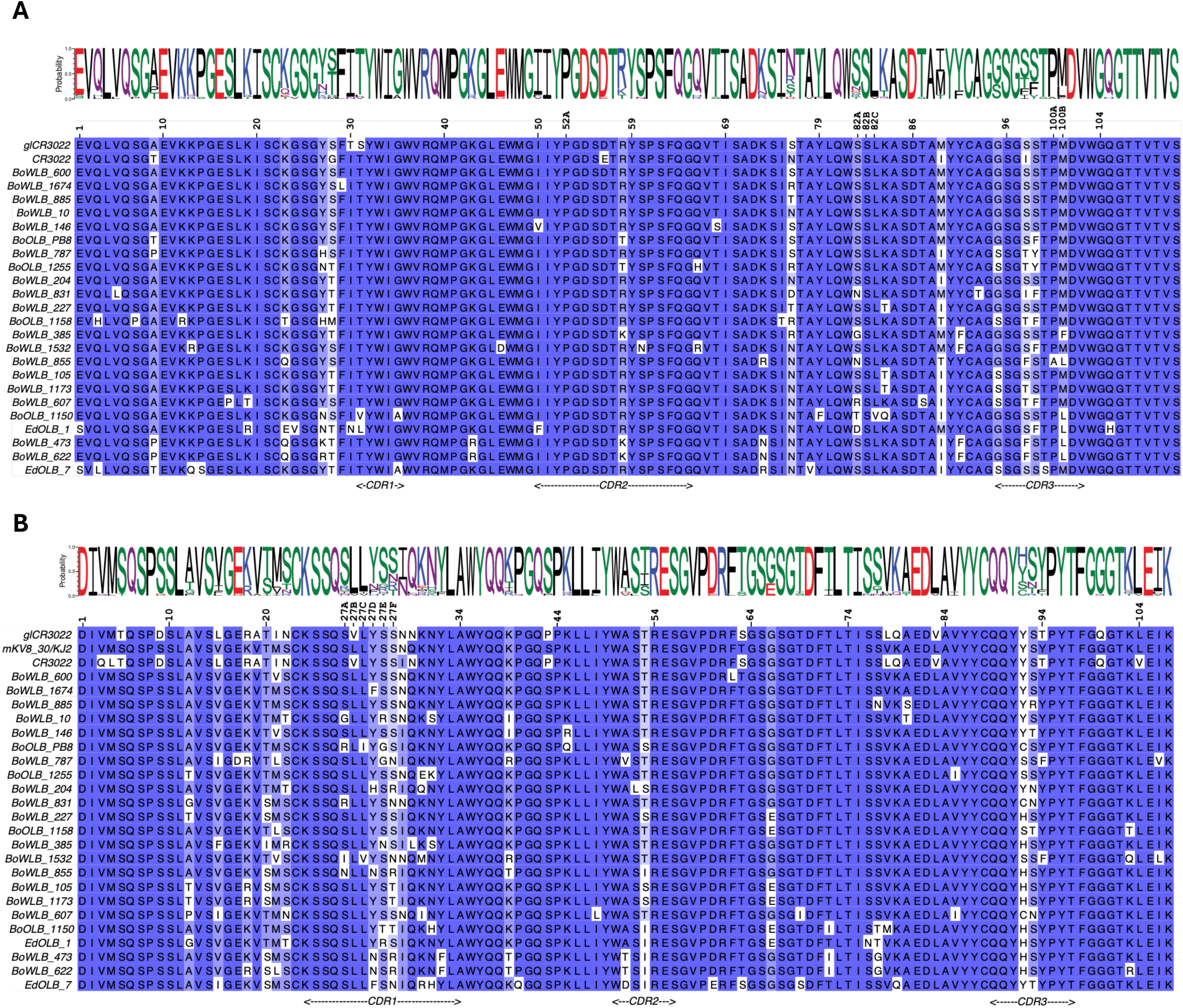
Multiple sequence alignments for. **(A)** heavy chains and **(B)** light chains of 23 mAbs, together with germline and affinity-matured CR3022 and the mouse germline *Igkv8-30*/*Igkj2* as references. The antibody numbering and CDRs delimiting (both based on Kabat) are denoted on the top and bottom of the alignment blocks, respectively. The Weblogo plots were generated after excluding the germline reference sequences.

**Supplementary Figure 5.**
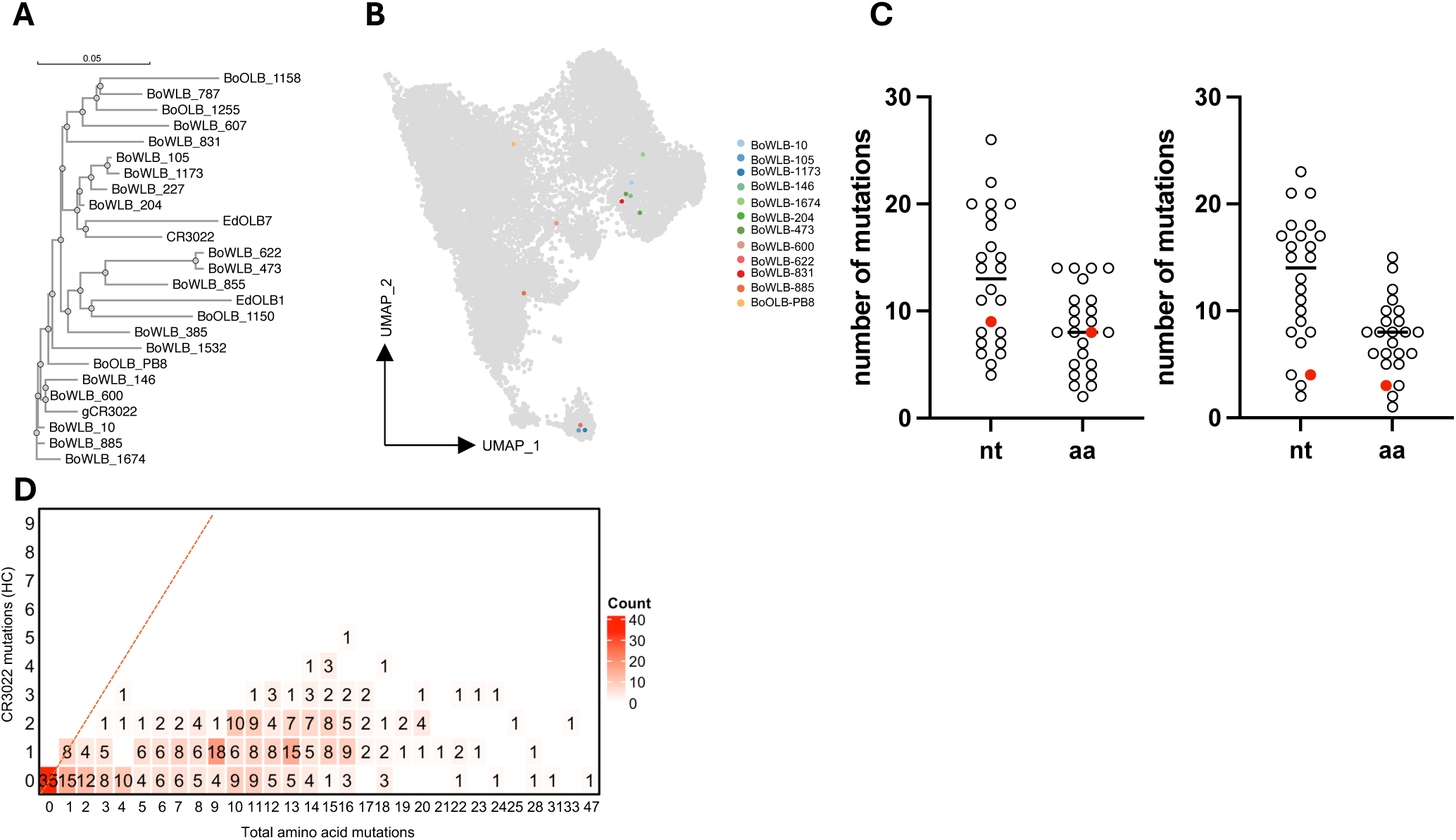
(**A**) Phylogenetic tree of expressed 23 mAbs including CR3022; **(B)** The distribution of a portion of expressed BCR clones among cluters **(C)** The number of HC and LC nucleotide and amino-acid mutations for 23 expressed mAbs, the red dots represent the number of mutations for CR3022; **(D)** CR3022 mutations detected in the human *IGHV5-51* gene cDNA sequences with various specificities. The y-axis indicates the number of CR3022 mutations observed in a given *IGHV5-51* gene. The number within each square on the plot represents the number of *IGHV5-51* sequences that contained those specific numbers of CR3022-type mutations. The x-axis indicates the number of mutations that a set of *IGHV5-51* sequences accumulated

**Supplementary Figure 6.**
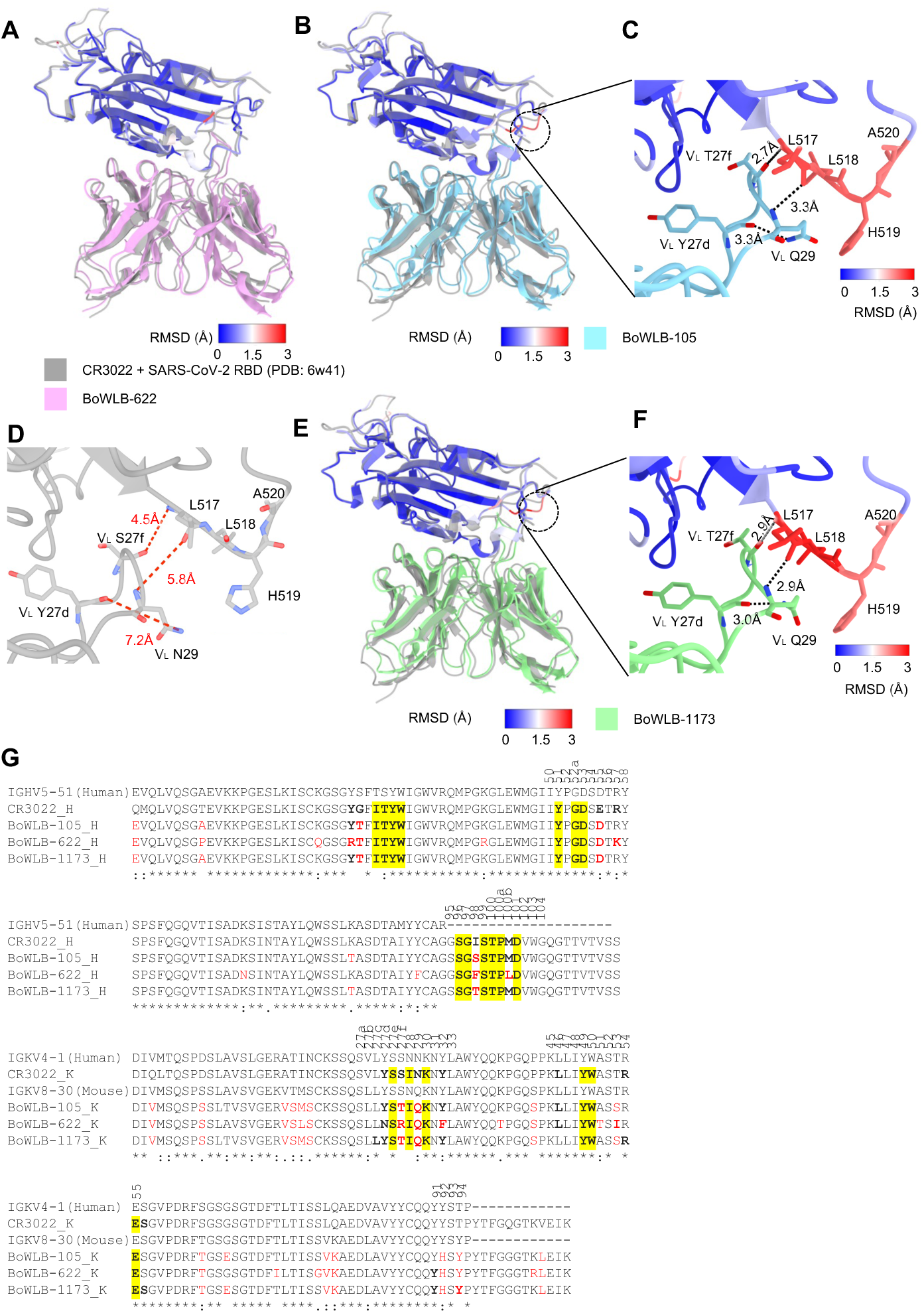
Structural superimposition and sequence alignment of prototype and the three engineered CR3022 antibodies. Kabat numbering is applied to the antibodies. Hydrogen bonds are represented by black dashed lines. CR3022 is colored grey. BoWLB-105, BoWLB-622, and BoWLB-1173 are colored cyan, pink, and green, respectively. Structural differences of RBD are color-coded by their root mean square deviation (RMSD). (A-C) Superimposition of the structures of RBD-CR3022 with, RBD-BoWLB-622, RBD-BoWLB-105, and RBD-BoWLB-1173, respectively. (D-F) Detailed interactions of RBD loop 517-520 with CR3022, BoWLB-105, and BoWLB-1173, respectively. Distances between atoms that form H-bonds are labeled. The red dashed lines and labels indicate the distance is too long to form an H-bond. (G) Antibody residues that are involved in interacting with the RBD (BSA > 0 Å^2^ as calculated by PISA ^78^) are in bold. Conserved paratope residues that are identical in all antibodies are highlighted by yellow boxes. Putative germline sequences were conducted by searching the IMGT database with IgBLAST ^80^. Residues identical in all aligned sequences are labeled by an asterisk (*), whereas a colon (:) and a period (.) indicate strongly similar and less similar sequences, respectively. Differences from CR3022 are shown in red. The sequence alignment was performed with Clustal Omega ^81^. Residue numbers of the CDR loops (Kabat numbering) are shown on top of the sequences.

**Supplementary Figure 7.**
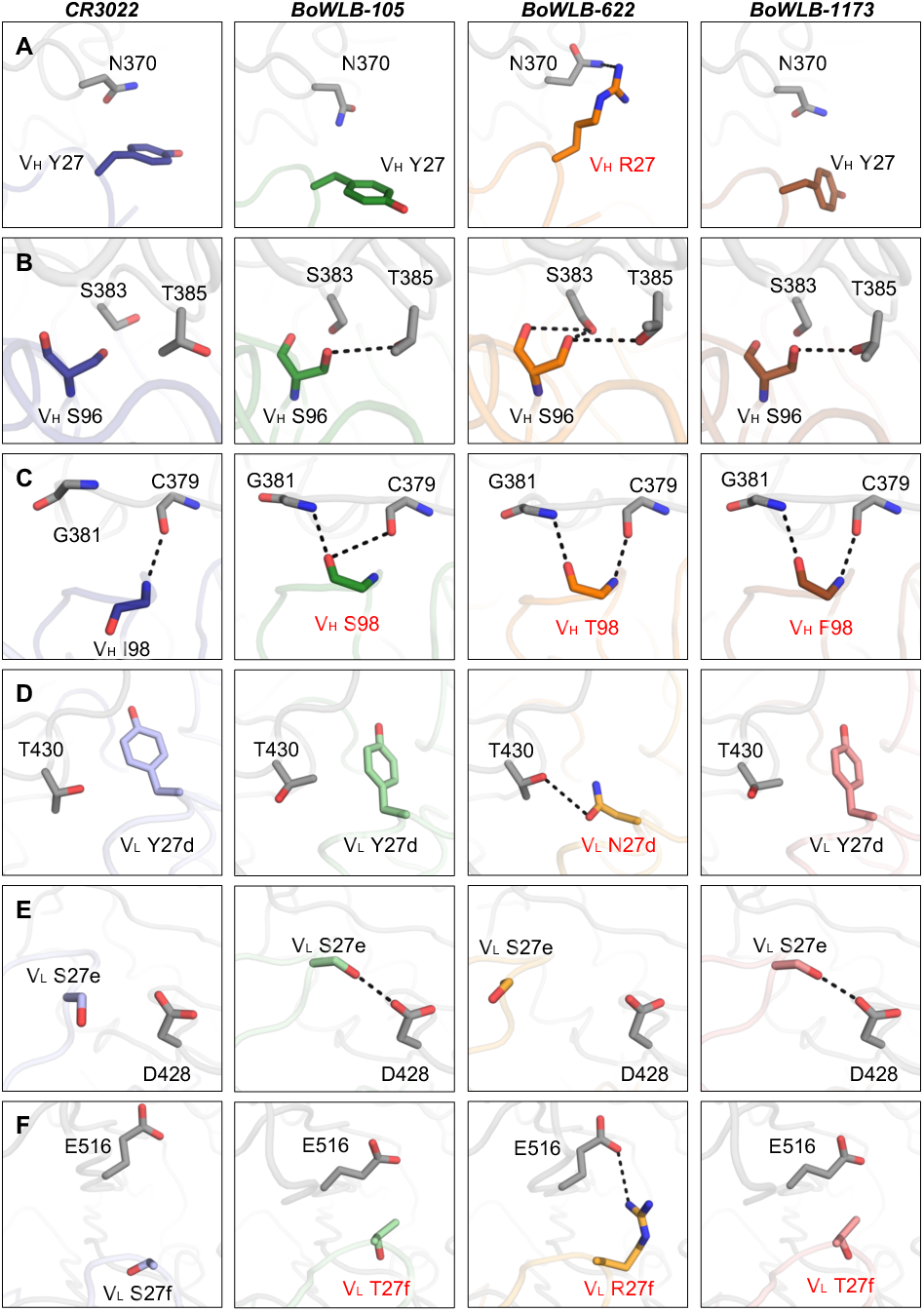
Structural comparison of single mutations and loop shift caused interaction difference between prototype and three engineered CR3022 antibodies in complex with SARS-CoV-2 RBD. Kabat numbering is applied to the antibodies. Hydrogen bonds and salt bridges are represented by black dashed lines. The color scheme of antibody-RBD complexes is the same as in Figure 6. (**A**) Comparison between the paratope V_H_ Y27 in CR3022, BoWLB-105, and BoWLB-1173 with its counterpart V_H_ R27 in BoWLB-622. V_H_ R27 forms H-bond with N370. (**B**) Comparison of the loop shift caused H-bond difference between V_H_ S96 and RBD S383 and T385. V_H_ S96 in BoWLB-105 and BoWLB-1173 form H-bonds with T385; V_H_ S96 in BoWLB-622 forms H-bonds with S383 and T385; while V_H_ S96 in CR3022 does not form any H-bond with S383 or T385. (**C**) Comparison of the main-chain conformation in heavy chain position 98. V_H_ I98 main-chain in CR3022 only forms interaction with C379 main-chain. However, main-chain V_H_ S98, T98, and F98 in BoWLB-105, BoWLB-622, and BoWLB-1173 form H-bonds with both C379 and G381main-chain. (**D**) Comparison between the paratope V_L_ Y27d in CR3022, BoWLB-105, and BoWLB-1173 with its counterpart V_L_ N27d in BoWLB-622. V_L_ N27d forms an H-bond with T430. (**E**) Comparison of the loop shift caused H-bond difference between V_L_ S27e and RBD D428. V_L_ S27e in BoWLB-105 and BoWLB-1173 form H-bonds with D428, while V_L_ S27e in CR3022 and BoWLB-622 do not form H-bond with D428. (**F**) Comparison between the paratope V_L_ S27f in CR3022, V_L_ T27f in BoWLB-105 and BoWLB-1173, and its counterpart V_L_ R27f in BoWLB-622. V_L_ R27f forms a salt bridge with E516.

**Table S4.**
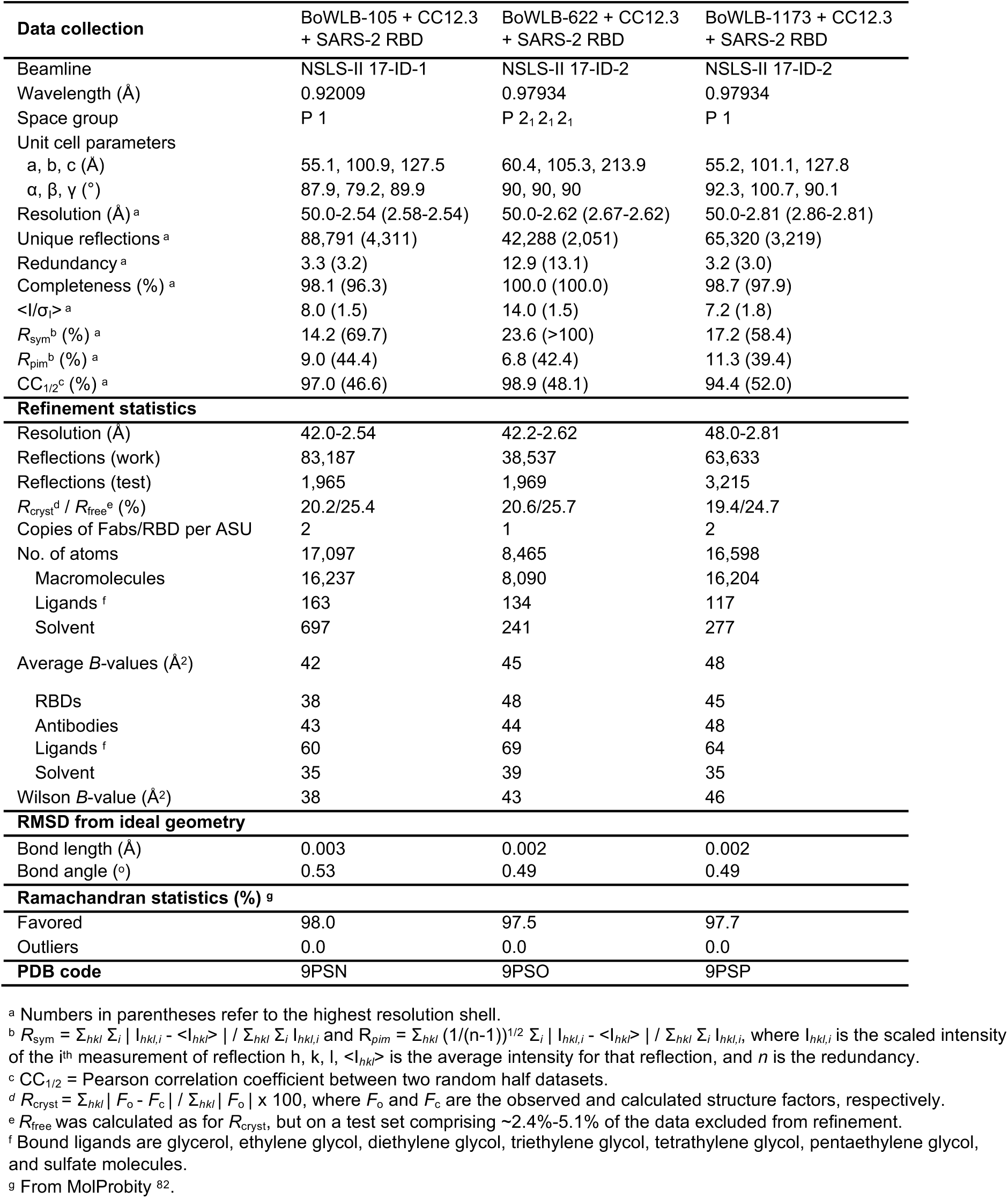
X-ray data collection and refinement statistics.

